# Functional classification of noncoding RNAs associated with distinct histone modifications by PIRCh-seq

**DOI:** 10.1101/667881

**Authors:** Jingwen Fang, Qing Ma, Ci Chu, Beibei Huang, Lingjie Li, Pengfei Cai, Pedro J. Batista, Karen Erisse Martin Tolentino, Jin Xu, Rui Li, Pengcheng Du, Kun Qu, Howard Y. Chang

## Abstract

Many long noncoding RNAs (lncRNAs) regulate gene transcription through binding to histone modification complexes. Therefore, a comprehensive study of nuclear RNAs in a histone modification-specific manner is critical to understand their regulatory mechanisms. Here we develop a method named Profiling Interacting RNAs on Chromatin by deep sequencing (PIRCh-seq), in which we profile chromatin-associated transcriptome in 5 different cell types using antibodies recognizing histone H3 and 6 distinct histone modifications associated with active or repressive chromatin states. PIRCh-seq identified chromatin-associated RNAs with substantially less contamination by nascent transcripts, as compared to existing methods. We classified chromatin-enriched lncRNAs into 6 functional groups based on the patterns of their association with specific histone modifications. LncRNAs were enriched with different chromatin modifications in different cell types, suggesting lncRNAs’ regulation may also be cell type-specific. By integrating profiles of RNA secondary structure and RNA m^6^A modification, we found that RNA bases which bind to chromatin tend to be more single stranded. We discovered hundreds of allele-specific RNA-chromatin interactions, nominating specific single nucleotide variants that alter RNA association with chromatin. These results provide a unique resource to globally study the functions of chromatin-associated lncRNAs and elucidate the basic mechanisms of chromatin-RNA interaction.

## INTRODUCTION

RNAs are both the product of transcription and major regulators of the transcriptional process. In particular, long noncoding RNAs (lncRNAs) are numerous in eukaryotes and function in many cases as transcription regulators^1–3^. With the development of next-generation sequencing (NGS), tens of thousands of lncRNAs have been revealed in both murine and human genomes, and have emerged as important regulators for different biological processes^4, 5^. However, among all expressed lncRNAs, only a small subset are shown to be cell essential^6^ or important for development^7^ or immune responses^8^. Strategies to annotate biochemical properties of lncRNAs will be helpful to prioritize lncRNA candidates for functional analyses. Some well-studied cases have indicated that one major mechanism of lncRNAs is their ability to function through binding to histone-modifying complexes^9, 10^. LncRNAs can either recruit chromatin modifiers to regulate the chromatin states or directly regulate the process of transcription through chromosome looping to bridge distal enhancer elements to promoters^11, 12^. Thereby, a genome-wide identification of chromatin-associated lncRNAs may reveal functions and mechanisms of lncRNAs in mediating chromatin modification and regulating gene transcription.

A considerable amount of literature has been published concerning protein-RNA interactions. The advent of technologies such as RIP^13^, CLIP^14^ and fRIP^15^ have led to the discovery of multiple protein-associated RNAs, including many chromatin regulators. Conversely, nuclear extraction methods followed by RNA-seq have enabled the detection of lncRNAs which are physically associated with chromatin^16–18^. In addition, more recently reported methods like GRID-seq^19^, MARGI^20^, and SPRITE^21^ can be used to capture pair-wise RNA interactions with DNA. However, these approaches are not capable of revealing which chromatin modifications are associated with specific lncRNAs, and are thus limited in the ability to elucidate their potential regulatory functions. For instance, a large number of lncRNAs are associated with Polycomb Repressive Complex 2 (PRC2), a key mammalian epigenetic regulator, to silence gene transcription by targeting its genomic loci and trimethylating histone H3 lysine 27 (H3K27me3)^22^. Therefore, lncRNAs associated with PRC2 complex may be enriched on heterochromatin regions with H3K27me3 modification. On the other hand, a new class of lncRNAs called super-lncRNAs were recently characterized. These lncRNAs target super-enhancers which have potential to regulate enhancer activities and transcription^23^. These super-lncRNAs may be enriched on euchromatin and active DNA regulatory elements with histone H3 lysine 27 acetylation (H3K27ac), H3 lysine 4 monomethylation (H3K4me1) and trimethylation (H3K4me3). Therefore, we believe it will be helpful to develop an experimental technology to distinguish different histone modification-associated lncRNAs, as well as analytical approaches to classify them and predict lncRNA functions based on their chromatin association patterns. Another technical challenge in studying chromatin associated lncRNAs is avoiding interference from abundant nascent transcripts on chromatin. For example, results from GRID-seq^19^ or MARGI^20^, approaches recently developed to identify in situ global RNA interactions with DNA, contain significant amounts of nascent transcripts, making it difficult to distinguish whether the detected RNA is truly chromatin-associated or merely captured during the process of transcription.

To address these questions, we developed a new method named Profiling Interacting RNAs on Chromatin followed by deep sequencing (PIRCh-seq), which enriches chromatin associated RNAs in a histone modification-specific manner and classifies functional lncRNAs based on the patterns of their attachment to nucleosomes with specific chemical modifications. Compared to current techniques for detecting chromatin-RNA association, PIRCh-seq efficiently reduces the influence of nascent transcripts with a significantly lower number of intronic reads. Through performing PIRCh-seq with histone H3 and a number of different histone modification antibodies on different cell types, we identified cell type-specific relationships between lncRNAs and epigenetics. We found that chromatin-associated lncRNAs can be classified into 6 functional groups based on their association with chromatin modifications, which undergo dynamic changes with cell differentiation. In addition, we found that bases on lncRNAs attached to chromatin tend to be more single stranded in an allele-specific manner. Overall, our PIRCh-seq data provides novel insights into global functional and mechanistic studies of chromatin-associated lncRNAs.

## RESULTS

### PIRCh-seq identifies RNA association with specific histone modifications in living cells

We conceived of PIRCh-seq as the inverse of ChIRP, a previously developed and robust method to crosslink endogenous RNA-chromatin interactions in living cells^24^. In the PIRCh-seq work flow, living cells are chemically crosslinked by glutaraldehyde and quenched with glycine, which prevents chromatin-associated RNA from further degradation. Chromatin is extracted and sonicated to 300-2000 basepair (bp) size, and then immunoprecipitated (IP) by histone modification-specific antibodies. Residual DNA and proteins are removed, and retrieved RNAs are then subjected to deep sequencing (**Figure 1A**). We tested the possibility that glutaraldehyde crosslinking may alter the pull-down specificity of antibodies targeting histone modification. Using SNAP-ChIP^25^, a pool of modified mono-nucleosomes with known histone tail modifications individually tagged with DNA barcodes, we found that glutaraldehyde crosslinking did not affect antibody specificity (**Figure S1A-C**). The input control for PIRCh is the lysate obtained after crosslinking and sonication but not subject to IP, which we also analyzed deep sequencing. RNAs that are retrieved by a histone modification over input beyond that expected by chance are considered PIRCh-seq hits. In this study, we generated and analyzed 26 high-resolution PIRCh-seq datasets from 2 different species: human and mouse; 5 cell types: human H9 embryonic stem cells (H9), human female fibroblasts (HFF), mouse V6.5 embryonic stem cells (mESC), mouse embryonic fibroblasts (MEF), and mouse neuronal precursor cells (NPC), targeting histone H3 and 6 histone modifications (namely H3K4me1, H3K4me3, H3K27ac, H3K27me3, H3K9me3 and H4K16ac) and input as control with 2 replicates for each experiment (**Figure S1D**). The expression distributions of the input RNAs extracted from our study were similar to that of the RNAs from nuclear extraction, but differed from cytoplasmic and total RNA (data obtained from GSE57231 & GSE32916 in the same cell line) (**Figure S1E**), suggesting that our chosen input could serve as a reasonable baseline for chromatin-associated RNA identification. Correlation analysis of these samples indicates the high reproducibility of PIRCh-seq experiments (*R*=0.900-0.988, **Figure S1F-M**).

**Figure 1.**
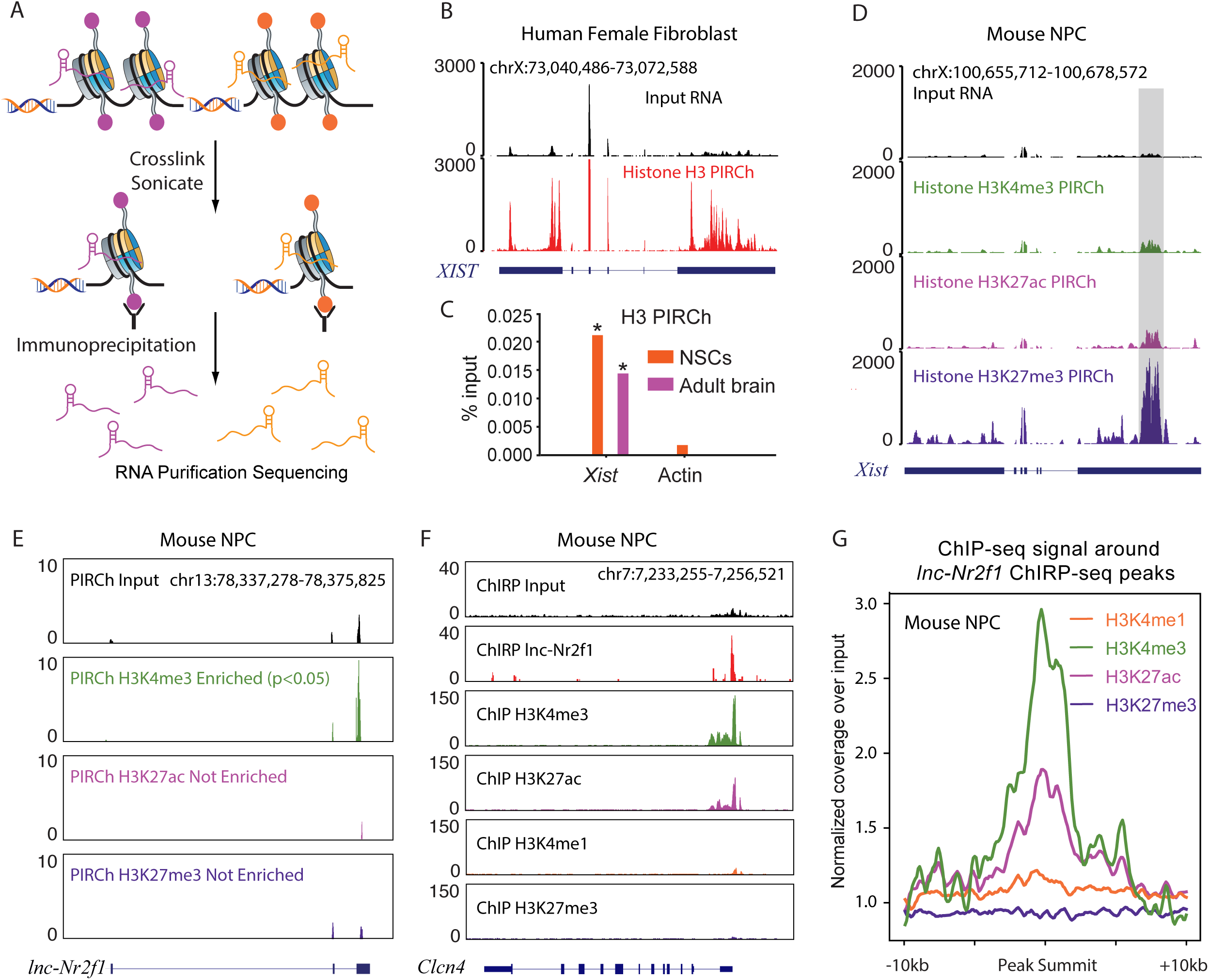
PIRCh-seq enables effective chromatin-RNA association *in vivo*. **A.** Schematic representation of PIRCh approach followed by high-throughput sequencing. **B.** Normalized input and histone H3 PIRCh-seq profiles of lncRNA *XIST* in human female fibroblasts. **C.** PIRCh-qPCR analysis in mouse neuronal stem cells (NSCs, orange) and adult brain (purple) shows that *Xist* is attached to chromatin H3 compared with Actin control. **D-E.** Normalized input and PIRCh-seq profiles with histone modifications of H3K4me3, H3K27ac, and H3K27me3 at the lncRNA *Xist* (**D**) and *lnc-Nr2f1* (**E**) locus in mouse neuronal precursor cells (NPC). **F.** Normalized input and ChIRP-seq profiles of lncRNA *lnc-Nr2f1* in NPC, and H3K4me3, H3K27ac, H3K4me1 and H3K27me3 ChIP-seq profiles in NPC. Showing *Clcn4* gene locus as an example. **G.** Average coverage of ChIP-seq signal (Reads per million over Input) around (+/-10kb) *lnc-Nr2f1* ChIRP-seq peaks in NPC.

As a proof of principle, we first examined PIRCh-seq signal of the well-characterized lncRNA *XIST*, which coats the inactive X chromosome in female cells, and is known to be associated with heterochromatin with repressive histone modifications^26^. Indeed, we observed that the PIRCh-seq signal of *XIST* is highly enriched on histone H3 over input in human female fibroblast cells (**Figure 1B**), as was histone H3 PIRCh followed by qRT-PCR for *Xist* in female murine neural stem cells (NSCs) and intact adult brain (**Figure 1C**). These results suggest that PIRCh-seq is not only capable of enriching chromatin-associated lncRNAs, but may be applied to study brain tissue *in vivo*. Similarly, the lncRNA *KCNQ1OT1*, which is involved in imprinting in Beckwith-Wiedemann syndrome by silencing lineage-specific transcription through chromatin regulation^27^, is also enriched on histone H3 over input, as expected (**Figure S2A**). Additionally, the imprinted oncofetal lncRNA *H19*^28^ was also enriched by histone H3 PIRCh-seq (**Figure S2B**). On the other hand, abundant protein-coding and house-keeping mRNAs, such as *ACTB* or *EEF2*, did not show PIRCh-seq enrichment as expected for cytoplasmic mRNAs (**Figure S2C-D**).

Next, we checked whether PIRCh-seq could enrich for RNAs associated with specific histone modifications. We performed PIRCh-seq on female NPCs with 3 ENCODE consortium validated antibodies targeting H3K4me3, H4K27ac and H3K27me3. PIRCh-seq in female NPCs demonstrated that *Xist* RNA was enriched by H3K27me3, a repressive mark enriched on the inactive X-chromosome, but not by active histone marks H3K4me3 nor H3K27ac that are depleted on the inactive X (**Figure 1D**). Interestingly, from *Xist’s* PIRCh-seq signal it is possible to infer which domain of this lncRNA is associated with chromatin. Within the *Xist* locus, the 5’ domain of *Xist* displays significantly more substantial enrichment in H3K27me3 PIRCh-seq as compared to other regions along the RNA (highlighted by the gray box, **Figure 1D**), consistent with previous findings that this is the domain potentially associated with chromatin (repC domain)^29–31^. Conversely, coding genes such as *Actb* and *Eef2* were not enriched on chromatin with the same set of modifications (**Figure S2E-F**). These results were obtained from 3 different cell lines in 2 species and indicate that PIRCh-seq is able to identify histone modification specific chromatin-associated lncRNAs transcriptome-wide.

PIRCh-seq can also be utilized to identify novel histone modification-specific chromatin-enriched lncRNAs. In our NPC PIRCh-seq, a lncRNA upstream of the *Nr2f1* gene, *lnc-Nr2f1*, was retrieved by the promoter marks histone H3K4me3 (*P*<0.05), but not enhancer-associated nor repressive modifications (H3K27ac and H3K27me3), indicating that this lncRNA may preferentially associate with H3K4me3 regions (**Figure 1E**). Recently, *lnc-Nr2f1* was reported to play a critical role in regulating neurodevelopmental disorders^32^. In order to further validate the chromatin-RNA association of this lncRNA, we retrieved *lnc-Nr2f1* RNA and mapped its associated DNA elements in NPCs (ChIRP-seq experiment). Overlaying *lnc-Nr2f1* ChIRP-seq with ChIP-seq data of the histone modifications confirmed that *lnc-Nr2f1* does bind to genomic locations with H3K4me3 (**Figure 1F-G**), further confirms that the PIRCh approach can retrieve lncRNAs specifically associated with certain modifications. In addition, gene ontology analysis of *lnc-Nr2f1* ChIRP-seq peaks using GREAT^33^ suggests that *lnc-Nr2f1* regulates cerebellar cortex development (**Figure S2G**, *P*<10^-5^), consistent with previous findings regarding the function of this lncRNA. These results not only demonstrate the reliability of PIRCh-seq in identifying chromatin-associated ncRNAs, but also suggest potential application of the histone modification-specific PIRCh-seq approach in predicting their functions.

### PIRCh-seq enriches lncRNAs on chromatin with low nascent transcription

Various techniques have been developed to study ncRNA functions on chromatin. For instance, ChIRP^24^, CHART^34^, and RAP^35^ are RNA-centric methods that profile DNA binding sites genome-wide of one target RNA at a time. Many investigators have isolated chromatin-associated RNAs from stringent nuclear or chromatin fractionation^16, 18^. In addition, recent methods such as GRID-seq and MARGI can be applied in mapping the global RNA-chromatin interactome^19, 20^. Comparatively, chromatin fractionation and sequencing detects chromatin-associated RNA without delineating the specific chromatin states that specific RNAs prefer. Furthermore, proximity ligation methods predominantly detect nascent RNAs co-transcriptionally tethered to chromatin by RNA polymerase, confounding signals from the functional chromatin-associated ncRNAs and background signal from all RNAs in the process of transcription. Thus, to evaluate the level of nascent transcription from PIRCh, we compared our PIRCh-seq results in H9 and HFF with that from GRID-seq^19^, che-RNA isolation (named CPE “chromatin pellet extract” for experiment and SNE “soluble-nuclear extract” for background control)^18^, and chromatin-associated RNAs (CAR)^16^. These experiments were all performed in human cell lines. We found that the ratios of intronic reads in PIRCh-seq profiles were significantly lower than those from previously reported methods (*P*<0.01, T-test), and were almost comparable with input RNAseq from bulk cultured cells (**Figure 2A**). Moreover, by averaging signals over the entire transcriptome centered by introns from all the existing methods, we found PIRCh was more effective in obtaining mature RNAs than extant chromatin-RNA enrichment methods, based on the higher signal over exons than introns (**Figure 2B**). We obtained similar findings in other cell types and with every tested histone modifications (**Figure 2C**, PIRCh-seq of V6.5 mouse ES cells with histone H3 and 6 histone modifications). These results demonstrate that PIRCh-seq consistently generates a significantly lower level of intronic reads with multiple histone modifications than existing methods, and therefore is able to preserve regulatory interactions in trans between lncRNAs and chromatin.

**Figure 2.**
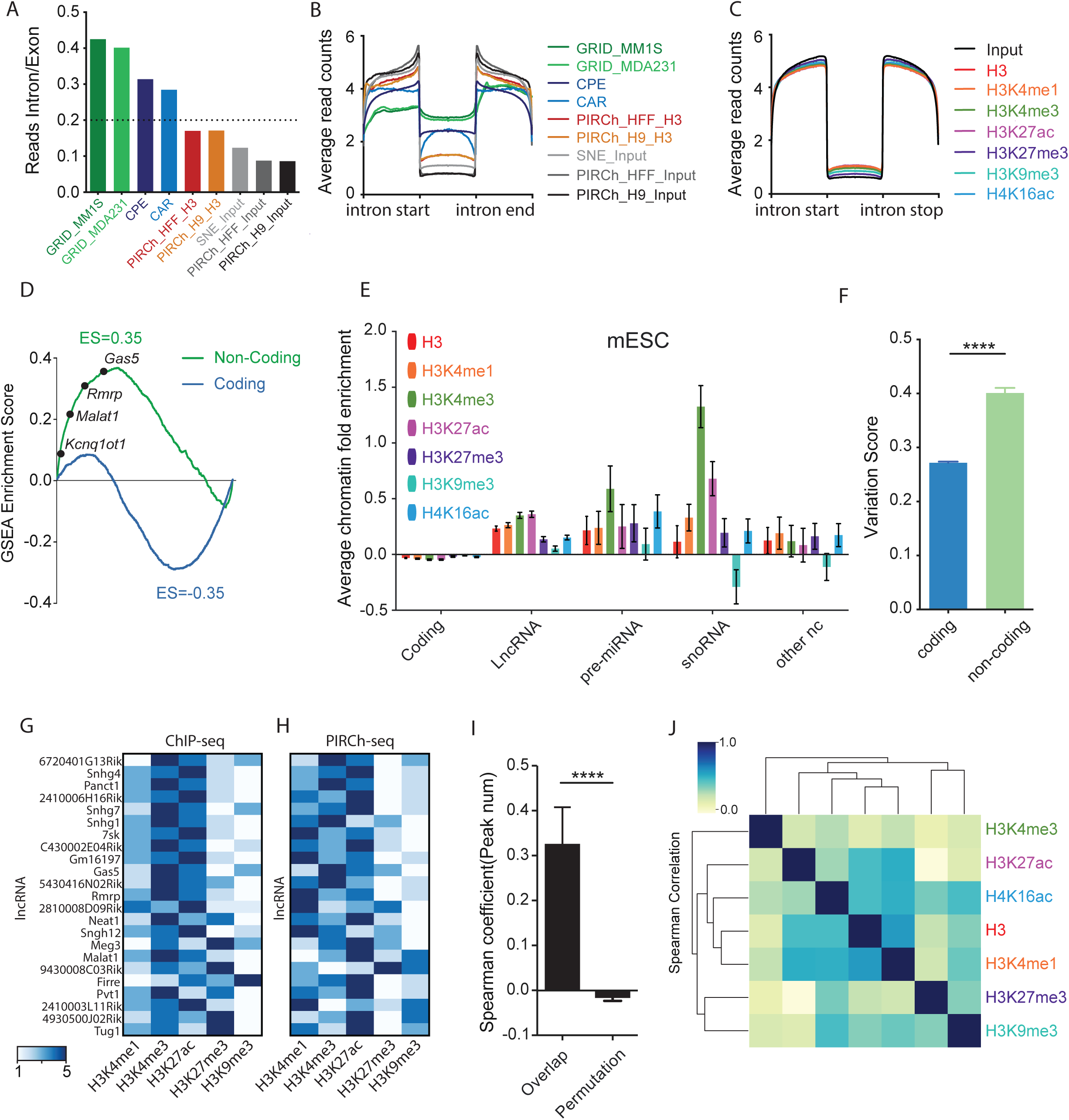
ncRNAs are enriched on chromatin compare with protein coding transcripts. **A.** Ratio of intronic over exonic reads obtained from different chromatin-RNA association sequencing technologies (GRID, CPE, CAR, and PIRCh) versus input controls in multiple cell lines. **B.** Normalized average read coverage around introns from different chromatin-RNA association sequencing technologies (GRID, CPE, CAR, and PIRCh) versus input controls in multiple cell lines. **C.** Normalized average read coverage around introns from histone modification specific PIRCh-seq profiles (colored) and inputs (black) in mouse embryonic stem cells (mESCs). **D.** Gene set enrichment analysis (GSEA) shows highly statistical enriched (FDR=0, P<0.0001) of non-coding genes (Green) and depleted of coding genes (Blue) on histone H3 in mESCs. Genes were ranked by their histone H3 PIRCh enrichment scores. **E.** Average fold enrichment (calculated by limma in R) of the coding gene, lncRNA, pre-miRNA, snoRNA and other ncRNA from histone modification specific PIRCh-seq profiles (namely H3, H3K4me1, H3K4me3, H3K27ac, H3K27me3, H3K9me3, and H4K16ac) in mESC. Error bar shows the standard deviation from the mean. **F.** Average variation score of the PIRCh-seq signals for the coding versus non-coding genes (****P<0.0001, two-tailed Welch’s T-test). Error bar shows the standard deviation from the mean. **G.** Heatmap displaying the ranking of the ChIP-seq enrichment of the chromatin binding sites of 23 lncRNA. The 23 lncRNAs are chromatin enriched from PIRCh-seq and the chromatin binding sites are obtained from ChIRP/CHART/RAP/GRID-seq profiles from the LnChrom database. Colors represent ranking from 1-5. **H.** Heatmap shows the ranking of PIRCh-seq enrichment of the same lncRNAs in **G**. Colors represent ranking from 1-5. **I.** Bar plot of the Spearman correlation coefficients between the ranking in **G** and **H** for each lncRNA versus random permutation (****P<0.0001, two-tailed Welch’s T-test). **J.** Unsupervised clustering of the Pearson correlation coefficients matrix of the histone modification specific PIRCh-seq profiles based on the enrichment scores from the 258 chromatin associated ncRNAs in mESC.

To further estimate the level of nascent transcription, we then integrated each histone modification-specific PIRCh-seq profile with its corresponding ChIP-seq dataset in the same cell line (V6.5 mESCs), and asked whether the PIRCh-seq signal of each RNA correlated with the nearby ChIP-seq signal carrying the corresponding modification (see **Methods**). The ChIP-seq profiles of each histone modification in mESC were obtained from ENCODE. Our results suggest that there was no significant correlation with these two sets of signals (**Figure S3A-E**), confirming that the nascent transcription from PIRCh-seq is negligible. These results suggest that the majority of PIRCh-seq enriched chromatin-associated RNAs are mature RNAs with introns spliced out, which allows PIRCh-seq to identify more chromatin-associated RNAs with low abundance, such as many ncRNAs.

### PIRCh-seq identifies ncRNAs associated with specific histone modifications

Because PIRCh-seq enables transcriptome-wide annotation of chromatin-RNA association, we next determined whether various types of RNA (especially coding RNAs versus ncRNAs) are differentially affiliated with chromatin. We first applied the limma package^36^ in R to normalize the RNA read counts in PIRCh-seq and input samples (**Figure S4A-B**). We then defined a PIRCh enrichment score by dividing the normalized read counts in PIRCh over input, and ranked all the transcripts by their enrichment scores in H3 PIRCh-seq. To test whether ncRNAs were enriched on chromatin, we performed a gene set enrichment analysis (GSEA)^37^ of the annotated coding and noncoding RNAs. We found that ncRNAs, but not coding RNAs, were indeed highly enriched on chromatin and many known ncRNAs were top ranked in terms of chromatin enrichment scores (**Figure 2D**). Next, we performed PIRCh-seq with antibodies specific to distinct histone modifications in mESCs. Similar to the enrichment on histone H3, we expect that ncRNAs should be highly ranked by the average fold enrichment of the histone modification-specific PIRCh-seq signal versus the corresponding input, among all the expressed genes. Indeed, compared with mRNAs, we found that in most cases (22 out of 28) the average enrichment scores of the annotated lncRNAs, pre-miRNAs, snoRNAs, as well as other ncRNAs were significantly higher on H3 and multiple histone modified chromatin than coding genes (**Figure 2E**, *P*<0.05, T-test). We then checked the distributions of the expressed and chromatin associated RNAs on histone H3 and chromatin with other modifications in mESCs, and found that ncRNAs were significantly more frequent on chromatin compared with mRNAs (**Figure S4C**), serving as additional evidence that ncRNAs are more enriched on chromatin in general. Furthermore, when we defined a variation score which measured the standard deviation of the chromatin association enrichment scores across each histone modification for every expressed RNA, we concluded that ncRNAs are significantly more variable than mRNAs (**Figure 2F**, *P*<0.001, T-test). This suggests that non-coding transcripts are more differentially enriched at distinct chromatin states, consistent with the potential regulatory function divergence of lncRNAs, and naturally prioritizes downstream studies of lncRNAs by activity.

We then sought to characterize the ncRNAs significantly on chromatin in mESC from their PIRCh-seq profiles. We considered PIRCh-seq biological replicates versus the inputs in limma^36^ and defined an RNA with chromatin association by P-value<0.05 (**Methods**). Using this cutoff, we identified 258 chromatin associated ncRNAs in mESC which were enriched in at least one of the 6 histone modification-specific PIRCh-seq profiles (**Table S1**). To further evaluate the performance of the PIRCh approach, we compared our PIRCh-seq enriched lncRNA results with 96 published RNA-chromatin association profiles from ChIRP/CHART/RAP/GRID-seq datasets, collected by LnChrom^38^. We found a total of 23 lncRNA, including *Xist*, *Firre*, *Rmrp*, *Tug1* and etc. were also expressed in our mESCs. All 23 lncRNAs were positively enriched in PIRCh, and 14 were significant with *P*<0.05, reaffirming the sensitivity of the PIRCh approach in identifying chromatin associated lncRNAs. Furthermore, we wanted to validate whether the PIRCh lncRNA enrichment patterns were consistent with results obtained from published orthogonal methods. We hypothesized that if a lncRNA is able to associate with DNA elements marked by a specific histone modification, its genomic binding sites from ChIRP/CHART/RAP/GRID-seq experiments should greatly overlap with corresponding ChIP-seq peaks associated with the same modification. We then obtained the genomic binding sites (peaks) of the 23 lncRNAs from the aforementioned experiments, and found the ratio of this overlap from published data (**Figure 2G**) is highly correlated with the corresponding PIRCh-seq signal among most of the lncRNAs (**Figure 2H**). The Spearman correlation coefficients of the ratio of the overlap ChIP-seq peaks^39^ with the lncRNA’s PIRCh-seq enrichment scores in the same cell line were significantly higher than random permutations (**Figure 2I**, *P*<0.0001). These results further confirm that PIRCh-seq reliably identifies chromatin associated lncRNAs.

Conversely, we hypothesized that certain ncRNAs are enriched at chromatin with distinct types of DNA regulatory elements, and asked whether gene regulatory elements could be naturally differentiated via chromatin-ncRNA association. We then calculated the pairwise Pearson correlation of all chromatin states based on the PIRCh-seq enrichment scores of 258 chromatin associated ncRNAs. It is clear that the enhancer-like states (H3K27ac, H4K16ac, H3K4me1) clustered together, then the promoters (H3K4me3), while the repressive histone modifications (H3K27me3, H3K9me3) were grouped in a distinct cluster (**Figure 2J**). Interestingly, the PIRCh-seq signal of histone H3 clustered closest with H3K4me1 (Pearson correlation r=0.89). We observed that H3K4me1 ChIP-seq signal from the same cells as above covers three to four times the genomic regions than other chromatin modifications, which may reflect the differential sensitivities of the different antibodies for ChIP (**Figure S4D**).

### PIRCh-seq classifies functional ncRNAs via chromatin association

Different gene regulatory elements--such as enhancers, promoters, insulators, and silenced elements carry distinctive and characteristic histone and DNA modifications (**Figure 3A**)^40^. We noticed that 14-25 ncRNAs in HFF and H9 respectively were also reported as “essential” ncRNAs with functions through CRISPRi screening^6^. We then hypothesized that specific modification enriched ncRNAs regulate each of these elements, and thereby the functions of ncRNAs can be classified by their divergent chromatin modification enrichment. Hence, PIRCh-seq is anticipated to classify and associate ncRNAs with functions such as promoter, enhancer, silencer, or insulator etc. To test this hypothesis, we analyzed *7sk*, a well-known regulator of RNA polymerase II elongation that resides at enhancers, promoters, and super enhancers^41^, consistent with its role in enhancer-promoter interactions. From *7sk* ChIRP-seq data in mESC, we noticed that its chromatin occupancy sites greatly overlapped with ChIP-seq peaks of H3K4me1, H3K4me3, and H3K27ac in the same cell type (**Figure 3B**), confirming an active function of *7sk*. Consistently, PIRCh-seq signal of *7sk* in mESC was also enriched at chromatin carrying these three histone modifications, but depleted of repressive modifications such as H3K27me3 and H3K9me3 (**Figure 3C**), suggesting the possibility to extrapolate lncRNA function using PIRCh-seq.

**Figure 3.**
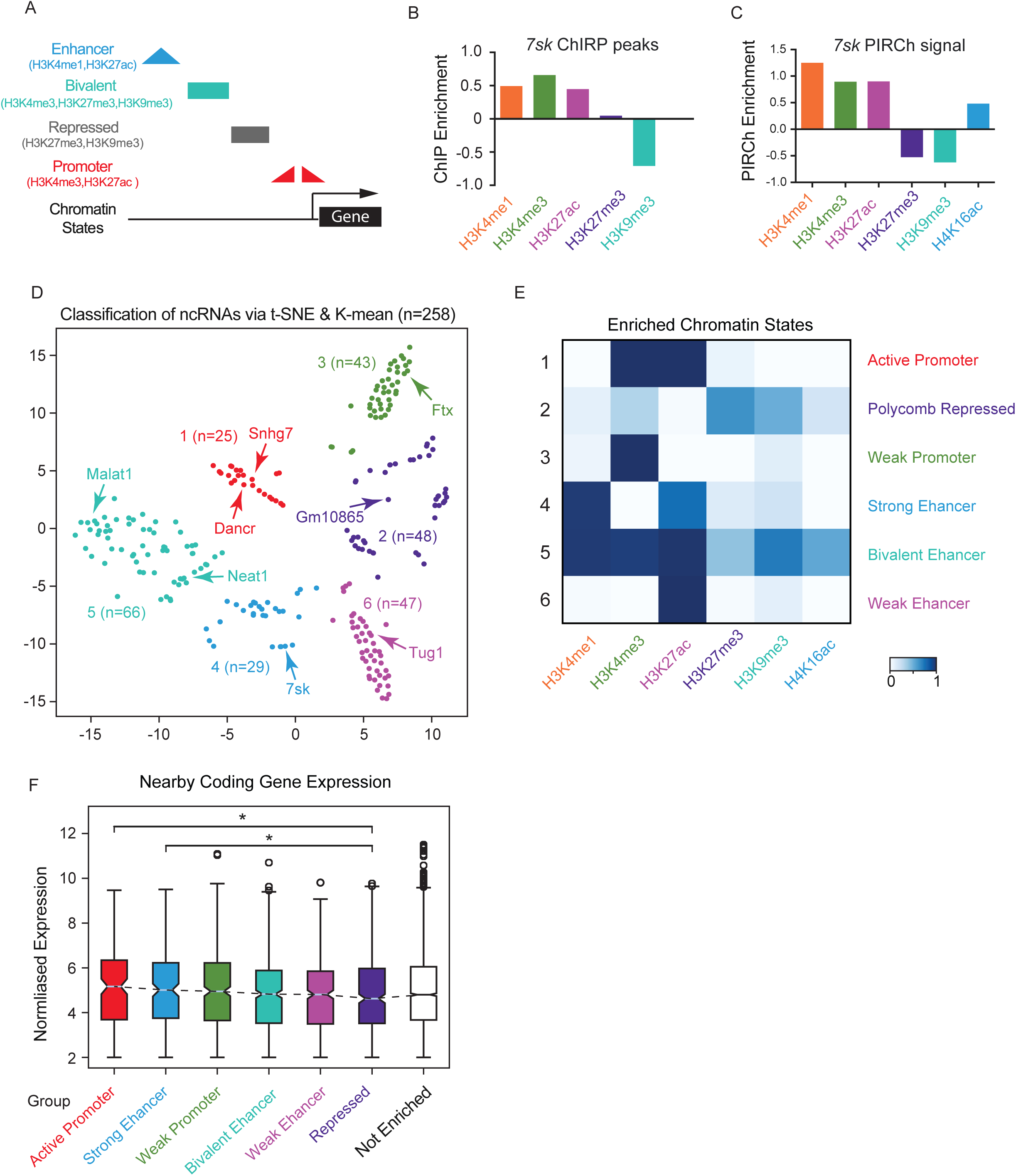
PIRCh-seq classifies functional ncRNAs via chromatin state association. **A.** Summary of histone modifications representing distinct regulatory patterns. **B.** The enrichment of the *7sk* ChIRP-seq peaks overlap with different histone modification ChIP-seq peaks in the same cell line (mESC). A positive value indicates the ChIRP-seq peaks are highly enriched with ChIP-seq peaks compare to random, and a negative value indicates depletion._ _ **C.** The PIRCh enrichment score of the lncRNA *7sk* in mESC from distinct histone modification specific PIRCh-seq experiments. A positive value means enriched, and a negative value means depleted. **D.** Classification of the PIRCh-seq identified chromatin associated ncRNAs (n=258) in mESC. Scatter plot shows the t-SNE result on PIRCh-seq enrichment score matrix and annotated by K-means clustering. **E.** Functional classification of histone specific chromatin-RNA association patterns defined by chromHMM algorithm. **F.** Box plot of the expression of the coding genes nearby (+/-100Kb) each group of PIRCh clustered ncRNAs defined in **D**. Center lines represent mean values; box limits represent the interquartile range; whiskers each extend 1.5 times the interquartile range; dots represent outliers. The expression of the coding genes that close to the ncRNAs in the “repressed” group is significantly lower than those in the “active promoter/enhancer” group (P<0.05, two-tailed Welch’s T-test). Genes close to un-enriched ncRNAs are shown as control.

We then analyzed all 258 PIRCh enriched ncRNAs and sought to categorize their functions based on their PIRCh-seq signals. We found that these ncRNAs associate with chromatin in a combinatorial pattern, similar to those observed in ChIP-seq performed on histone modifications (**Figure S5A**). H3K27ac, H3K4me3 and H3K4me1 were the top 3 most favored chromatin states that interacted with ncRNAs, consisting of 88% of the enriched ncRNAs in mESC. As we know from histone ChIP-seq, instead of each individual modification, a combinatorial pattern of multiple modifications better classifies the functions of DNA elements^42^. A machine learning strategy employing hidden Markov model, named chromHMM, which automatically learns the major combinatorial patterns, was applied successfully to classify DNA elements based on histone modifications^43, 44^. We then inquired if a similar strategy could be used to classify chromatin-associated ncRNAs and examine if the functions of these ncRNAs could be distinguished based on their association with histone modifications. To investigate this relationship transcriptome-wide, we started from a 258 by 6 matrix of enrichment scores in mESC, where each row was an enriched ncRNA as defined above, each column was a histone modification, and each element of the matrix represented the enrichment score of the corresponding ncRNA on the specific modified chromatin (**Methods**). We then applied K-means clustering on the matrix, where the number of Ks was determined by the Silouette method^45^. This analysis yielded 6 distinct groups of chromatin-associated ncRNAs, which were visualized in a 2-dimensional projection of t-distributed stochastic neighbor embedding (tSNE) (**Figure 3D**). Within these 258 chromatin associated ncRNAs, 247 are lncRNAs, and many well-studied ncRNAs, such as *7sk*^41^, *Neat1*^10^ and *Malat1*^46^, and *Dancr*^47^ naturally clustered into groups with distinct function. Interestingly, 14 lncRNAs were also reported to have a biological functional based on LncRNAdb^48^, and 8 out of 56 were predicted bivalent in mESCs (**Figure S5B**, odds-ratio=4.4, *P*<0.01, Chi-square test). In addition, for each cluster we evaluated the relative contributions of each histone modification based on the enrichment pattern of the chromatin associated RNAs, and defined the clustered states by active promoters, heterochromatin, weak promoter, strong enhancer, bivalent, and weak enhancer (**Figure 3E**). Overall, we partially recapitulated the chromatin classifications based on chromHMM algorithm to ChIP-seq profiles^43^. These results suggest that the chromatin association of ncRNAs can be used to classify ncRNAs that might have functional implications.

Although tens of thousands noncoding transcripts were discovered in the past few years, only a small portion that function through chromatin organization were consolidated. More recently, evidence has accumulated that indicates many lncRNAs may regulate gene expression in cis^49, 50^. Since the PIRCh approach cannot pinpoint the exact binding sites of chromatin-associated lncRNAs, it does not directly predict whether each lncRNA is functioning in cis or trans. Instead, PIRCh provides more information about the epigenetic function of the lncRNA, in context of the histone modifications it associates with. Our analysis suggests that chromatin-associated lncRNAs function both in trans and cis. For example, when we calculated the nearby (+/-100Kb) coding gene expression of the PIRCh clustered ncRNAs in **Figure 3D**, we observed that lncRNAs were monotonically decreasing from the more active to more repressive groups; additionally, the nearby coding gene expression of the “Active Promoter” and “Strong Enhancer” lncRNA groups were significantly higher than that of the group “Repressed” ncRNAs (**Figure 3F**, *P*<0.05, T-test). However, when the chromatin associated ncRNAs were grouped based on their enrichment with each histone modification, no significant expressional differences were observed from nearby coding gens. (**Figure S5C**), e.g. compared H3K27me3 vs H3K27ac. No similar trends were observed in the expression patterns of the ncRNAs themselves (**Figure S5D**). These results not only indicate that the chromatin-associated ncRNAs may function through a combinational pattern of the histone modifications instead of an individual modification, but also favors the argument that the chromatin associated ncRNAs may function in cis in general. Nevertheless, not all the lncRNAs enriched in our PIRCh experiment function in cis. When we integrated each histone modification specific PIRCh-seq profile with its corresponding ChIP-seq signal at the genomic loci of the chromatin-enriched ncRNAs, no statistical correlation was observed (**Figure S3**), suggest that some lncRNAs can function in trans.

### Cell type-specific chromatin association of ncRNAs

It is known that ncRNAs are differentially expressed in distinct cell types and perform specific cellular functions. Therefore, we sought to check whether the patterns of ncRNA-chromatin association diverge in distinct mouse cell types, and how these patterns contribute to their cell type-specific functions. We then performed PIRCh-seq on MEF cells and analyzed the profiles in an identical fashion to the mESC data. Similar to the mESC results, we observed that PIRCh-seq identified lncRNAs enriched on chromatin with low nascent transcription (**Figure S6A**), and non-coding transcripts were consistently more enriched on chromatin compared with protein coding gene in MEF cells (**Figure S6B-C**), validating these conclusions in distinct cell types. We then performed a similar enrichment analysis on MEF and NPC PIRCh-seq profiles and obtained 200 and 110 chromatin associated ncRNAs respectively (*P*<0.05). The combinatorial patterns of the MEF enriched ncRNAs are predominantly similar to those from mESC (**Figure S6D-E**). As a negative control, the IgG PIRCh was tested in tandem with the other chromatin modification PIRCh experiments performed in MEF. Differential analysis of PIRCh-groups over IgG control revealed that only 1 out of 200 PIRCh-enriched ncRNA over input was also enriched in IgG, evincing the high specificity of our method in identifying the chromatin associated ncRNAs (**Figure S6F**). In our analysis, a total of 458 chromatin enriched ncRNAs were identified in three cell types, 20 of which were enriched in all three cell types (**Figure 4A**). We then calculated the Pearson correlation coefficient matrix based on the enrichment scores of these 458 ncRNAs. Unsupervised clustering of this correlation matrix suggested that the cell type specificity was the dominant factor which determines ncRNA chromatin association (**Figure 4B**).

**Figure 4.**
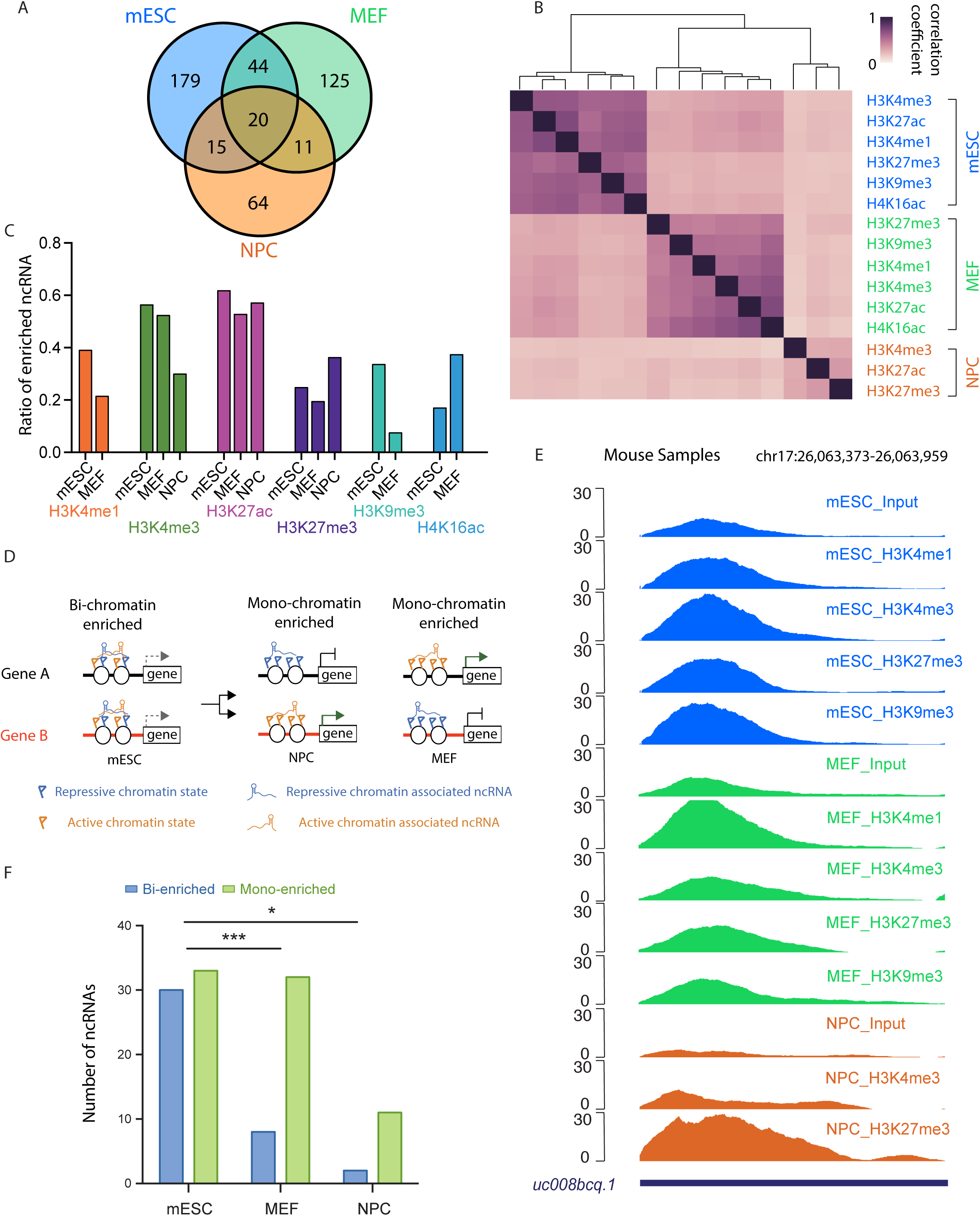
Cell type specific chromatin association of ncRNAs. **A.** Number of chromatin enriched RNAs in mESC, MEF and NPC. **B.** Unsupervised clustering of the Pearson correlation coefficients matrix of the histone modification specific PIRCh-seq profiles in mESC, MEF and NPC, based on the enrichment scores from the 458 chromatin associated ncRNAs in each cell type. **C.** Ratio of the chromatin enriched ncRNA under each chemical modification over the total number of enriched ncRNAs in mESC, MEF, and NPC. **D.** Schematic illustration of how RNAs enriched on both the repressive and active chromatin (bi-chromatin enriched) and either the repressive or active chromatin (mono-chromatin enriched). **E.** UCSC tracks of the normalized PIRCh-seq signal at the lncRNA *uc008bcq.1* locus in mESC, MEF and NPC. *uc008bcq.1* is bi-chromatin enriched in mESC, but mono-chromatin enriched in MEF and NPC. **F.** Number of ncRNAs that are bi-chromatin enriched or mono-chromatin enriched in mESC, MEF and NPC (***P<0.001, *P<0.05, Chi-square test).

Embryonic stem cells are characterized by their pluripotency - the ability to give rise to multiple cell types. The chromatin state in ES cells is reported to be more flexible than those of differentiated cells^51^. Interestingly, compared with those of the more differentiated cells (MEF and NPC), the ncRNA chromatin association in mESCs showed a higher correlation coefficient among distinct histone modifications, suggesting the specificity of chromatin-associated ncRNAs in mESCs is more plastic than those in differentiated cells (**Figure 4B**). In addition, we analyzed the percentage of enriched ncRNA versus total expressed ncRNA in each cell type for every tested chromatin modification, and found significantly more ncRNAs enriched on chromatin with H3K9me3 in ES cells when compared with MEF (**Figure 4C**, *P*<0.05, Chi-square test), but fewer on chromatin with H4K16ac. This result may reflect the joint presence of activating and repressive histone marks on genome regions, termed bivalent^52^ and trivalent chromatin domains^53^ in ES cells. We identified ncRNAs which were associated with both active and repressive histone marks consistent with bivalency, while others associated with strictly active or repressive marks (**Figure 4D**). Since PIRCh-seq enabled us to identify cell-type and histone modification specific ncRNA-chromatin associations, we first screened for ncRNAs which were enriched at both active and repressive chromatin in ES cells but only enriched in either active or repressive markers in differentiated cells. We found several ncRNAs of this description. For example, ncRNA *uc008bcq.1* is broadly enriched in ES cells with high PIRCh-seq signals associated H3K4me1, H3K4me3, H3K27me3 and H3K9me3 modifications, but enriched only on active chromatin of H3K4me1 in MEF and repressive chromatin of H3K27me3 in NPC, implying lineage-specific resolution of chromatin associations (**Figure 4E**). Interestingly, there were dozens of such ncRNAs that are distinctly enriched in certain cell types. Since ES cells possess a higher potential to differentiate into multiple lineages, and hence more poised chromatin states, we expected more bivalent-enriched (“bi-enriched” for short) and fewer mono-enriched ncRNAs in mESC compared with more differentiated cells such as MEF and NPC. In mESC, we found 30 bi-enriched and 33 mono-enriched ncRNAs; while in MEF, we found only 8 bi-enriched but 32 mono-enriched ncRNAs; lastly, in NPC, we found 2 bi-enriched and 11 mono-enriched ncRNAs (**Figure 4F**, *P*<0.01 for MEF and *P*<0.05 for NPC, Chi-square test). These results indicate that ncRNAs may play distinct functional roles by either enhancing or repressing gene expression or both in certain cell types, conducted by affixing to either active or repressive chromatin or both.

### Single-stranded RNA regions as candidate mediators of chromatin association

As a key player in the central dogma of biological regulation, RNA and its ability to adopt specific structures is intimately involved in every step of gene expression. Previously, multiple approaches have been described in order to probe RNA secondary structure transcriptome-wide *in vitro*^54^ and *in vivo*^31, 55^ in mammalian cells, revealing structural principles of RNA-protein interactions. Correspondingly, we noted that RNA enrichment on chromatin occurs in a domain-specific manner based on our PIRCh-seq data. For instance, the repC domain of *Xist* is dramatically more enriched on chromatin carrying H3K27me3 modifications (highlighted by the gray box, **Figure 1D**). *Malat1* is another well-studied chromatin-associated lncRNA which binds to active chromatin^10^. Instead of attaching to histone proteins across the entire transcript, we noticed from the H3K4me3 PIRCh-seq signal that there were certain regions on *Malat1* which were more closely associated with chromatin than the rest bases on the transcript (**Figure 5A**). Interestingly, these regions tend to be single-stranded according to both 2’ hydroxyl acylation profiling experiments (icSHAPE data) and RNA secondary structure predictions from RNAfold^56^ (**Figure 5B**). This led us to investigate whether there are structural preferences involved in RNA-chromatin association (**Figure 5C**). We first obtained a transcriptome-wide and per-base RNA secondary structure profile from icSHAPE data measured in mESCs^55^. A high icSHAPE score suggests a greater probability that a base is single stranded. We then applied a 5-base sliding window method to identify the enriched sites (peaks) on each RNA which interacted with chromatin from PIRCh-seq, compared with our input control (see **Methods**). We then overlaid the structural profiles from icSHAPE on top of all the histone modification specific PIRCh-seq peaks centered by the peak summits and generated an average structural profile for each modification. Our results show that bases ∼5-10 nt upstream of the chromatin associated peaks are more likely to be single stranded (**Figure 5D**). To test the significance of this single-strand preference, we performed 2-tailed Welch’s T-tests by comparing all the icSHAPE scores of the bases from PIRCh-seq peaks with those from a randomly selected background, and found this phenomenon was significant with *P*<10^-5^. We then asked whether RNAs containing a greater number of single-stranded bases are more likely to be associated with chromatin. We separated expressed RNAs into two groups based on chromatin enrichment or depletion, and calculated the average icSHAPE scores for every RNA in each group. We noticed that, on average, RNAs enriched on chromatin tended to be more single-stranded with higher icSHAPE scores (**Figure 5E**, *P*<0.001, T-test). Similarly, we took the top 100 most single-stranded RNAs and top 100 most double-stranded RNAs based on their average icSHAPE scores and confirmed that the average chromatin enrichment scores of the most single-stranded RNAs were significantly higher than those of the double-stranded RNAs (**Figure 5F**, *P*<0.01, T-test). These results suggest that RNAs containing more single stranded regions are more likely to associate with chromatin.

**Figure 5.**
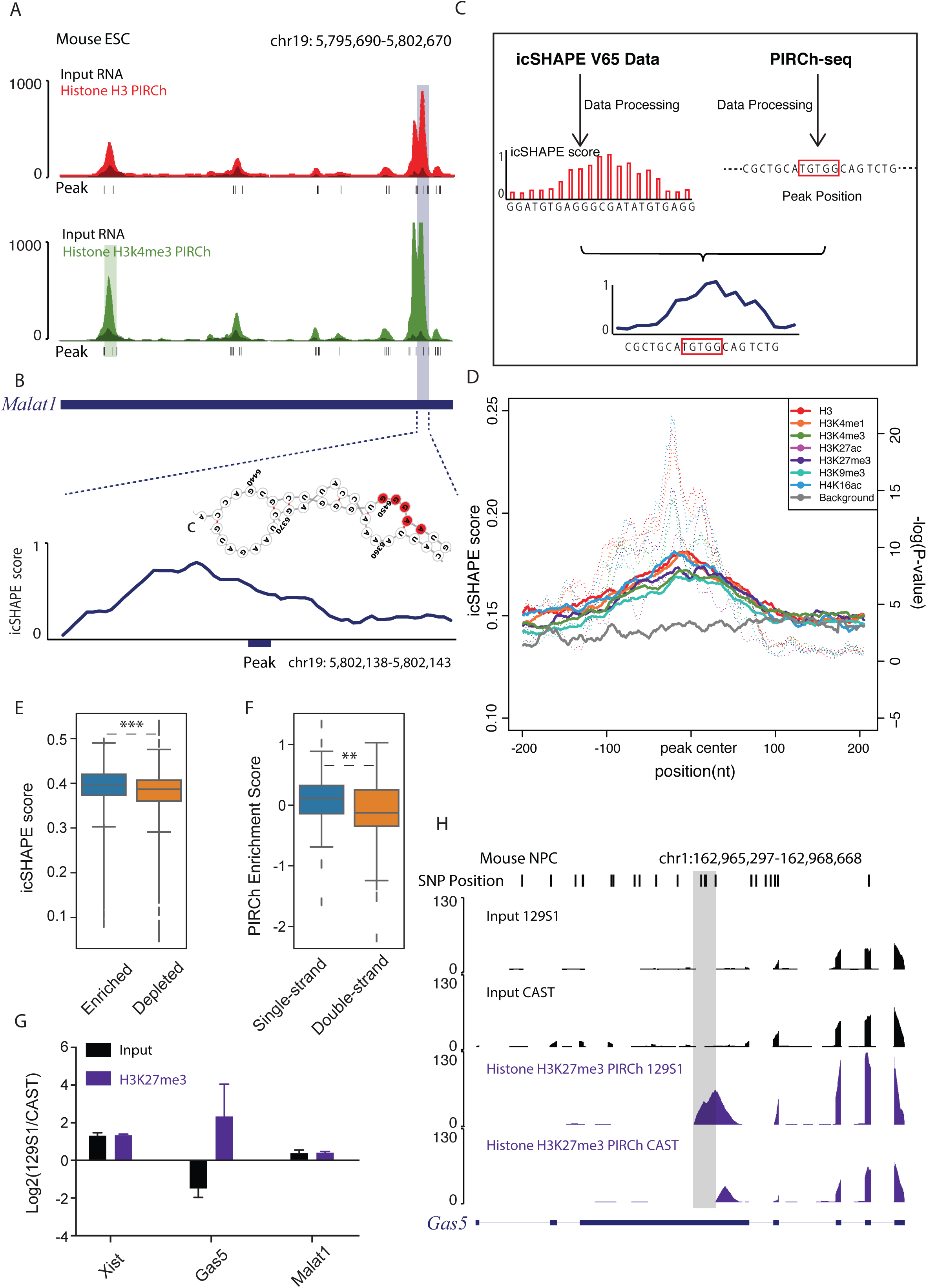
RNA with single strand are more likely to associate with chromatin. **A.** UCSC track of the normalized input (black) and H3 (red) and H3K4me3 (green) PIRCh-seq signals of lncRNA *Malat1* in mESC. Bottom peaks are chromatin enriched sites on *Malat1*. **B.** Structure profile from icSHAPE and structural prediction from RNAfold around a zoom in chromatin associated peak on lncRNA *Malat1*. **C.** Computational workflow to integrate RNA secondary structure information from icSHAPE and chromatin enrichment information from PIRCh-seq to study the structural preference of chromatin-RNA association. **D.** Average diagram of icSHAPE scores around all PIRCh-seq peaks under different histone modifications (colored solid line) versus a randomly selected background (grey solid line). P-values (colored dash line) were estimated by using two-tailed Welch’s T-test on every position between PIRCh-seq profiles over background. **E.** Box-plot of the icSHAPE score of PIRCh-seq enriched vs depleted RNAs (***P<0.001, two-tailed Welch’s T-test). Center lines represent mean values; box limits represent the interquartile range; whiskers each extend 1.5 times the interquartile range; dots represent outliers. **F.** Box-plot of the PIRCh-seq enrichment scores of the top 100 most single stranded RNAs versus the top 100 most double stranded RNAs based on icSHAPE scores (**P<0.01 two-tailed Welch’s T-test). Center lines represent mean values; box limits represent the interquartile range; whiskers each extend 1.5 times the interquartile range; dots represent outliers. **G.** Relative allele specific RNA expression and chromatin enrichment of lncRNAs *Xist*, *Gas5* and *Malat1* in the 129S1 allele versus the CAST allele of NPC. The 129S1 version of lncRNA *Xist* is highly expressed and also enriched at chromatin with H3K27me3 modification. Both alleles of lncRNA *Malat1* were almost equally expressed and enriched. The 129S1 version of *Gas5* was lowly expressed but highly enriched on chromatin compared to the CAST version of the same gene. **H.** Normalized allele specific input and histone H3K27me3 PIRCh-seq signals in the 129S1 and CAST alleles. Top shows single nucleotide polymorphisms (SNP) positions that distinguish the alleles.

### Single nucleotide variants and RNA modifications that alter chromatin association

Genetic variation can alter RNA structure and function *in vivo*. Single nucleotide polymorphisms (SNPs) comprise the most prevalent source of variation, and SNPs that alter RNA secondary structures, termed “riboSnitches” (a fusion of SNP and riboswitch), are a recently appreciated source of noncoding variants associated with human diseases^57^. We therefore asked whether different alleles of the same RNA may differentially associate with chromatin; and if so, how is it related to the RNA structure? In order to answer those questions, we performed PIRCh-seq in the NPC line that is derived from the F1 hybrid offspring of two mouse parental lines (129S1 and CAST) with a high density of SNPs across the genome (∼1 SNP per 100 nucleotides). We first built the reference genomes for each mouse line and aligned the raw reads to 129S1 and CAST separately with 0 mismatches to reduce false positive hits. Reads mapped to either 129S1 or CAST were counted to construct the allele-specific RNA expressions and chromatin enrichments profiles (see **Methods**). First, we looked at whether allelic RNA chromatin association is related to allelic expression. From allele-specific PIRCh-seq analysis, we found that for most RNAs, allelic or unbiased expression from the two alleles determines allelic or unbiased chromatin association pattern. For example, it is known that only the 129S1 version of the lncRNA *Xist* is expressed in this cell line. Consistent with allelic expression, we found that enrichment of *Xist* in H3K27me3 modification is much higher in 129S1 versus the CAST version of the lncRNA (**Figure 5G**). An additional example is the lncRNA *Malat1* in which both the 129S1 and CAST alleles are almost equally expressed. As predicted, we observed unbiased enrichment on chromatin for both alleles. Moreover, we discovered several lncRNAs that are enriched on chromatin in an allele-specific manner independent of the expression levels from the two alleles (**Table S2**). For example, *Gas5* is a lncRNA that binds to PRC2 complex and mediate transcriptional repression^58^. We found that *Gas5* is enriched in H3K27me3 modification, consistent with its understood repressive function. Notably, even though the CAST version of *Gas5* was 3-fold more expressed in the input sample, the 129S1 allele was 4-fold more enriched on chromatin carrying H3K27me3 modification (**Figure 5G**, *P*<0.05, T-test), suggesting that 129S1 allele of *Gas5* preferentially associates with chromatin. To further investigate the mechanism under *Gas5* allele-specific enrichments, we predicted the secondary structure of the 129S1 and CAST version of *Gas5* using RNAfold (**Figure S7A, B**), and found that several riboSnitches (1774 T/C, 1804 C/T, 1810 T/A, 1812 T/G, 1887 T/C, CAST(mm9)/129S1) converted one of the chromatin binding sites of *Gas5* from single-stranded in 129S1 to double-stranded in CAST and thus depleted its association with repressive chromatin in the latter (**Figure 5H**). Consistent with this prediction, when we calculated the icSHAPE score obtained from mESC containing 129S1 allele^59^ for the *Gas5* region allelic enriched in H3K27me3, we concluded that the region is more likely to be single stranded (**Figure S7C**).

Another major factor that can influence RNA structure is RNA modification, such as the *N6*-methyladenosine (m^6^A) modification. Previous studies have shown that m^6^A can alter base-pairing thermodynamics and destabilize RNA duplexes^55, 60, 61^. We also evaluated whether RNA modifications affect RNA-chromatin association. We integrated PIRCh-seq data with the transcriptome-wide profiles of RNA m^6^A modifications in mESCs from our previous study^62^, and found the distribution of PIRCh-seq peaks along the transcripts is similar to that of m^6^A modified regions (**Figure S8A**). When we overlaid m^6^A signals on top of PIRCh-seq peaks, we found that RNA bases associated with chromatin are generally more m^6^A modified (*P*<10^-5^ in H3, **Figure S8B-C**). These results may reflect that the tendency of m^6^A to induce RNA single-stranded regions that coincide with elements for chromatin association, or due to additional mechanisms that jointly impact chromatin association and RNA modification.

## DISCUSSION

### PIRCh-seq identifies chromatin associated RNAs genome-wide

A large and growing body of literature has investigated protein-RNA interactions. The development of approaches such as RIP^13^, CLIP^14^ and fRIP^15^ have enabled the successful elucidation of many RNAs associated with proteins, including multiple chromatin regulators. Studies have also shown that many lncRNAs function through DNA/chromatin interaction. Previously described techniques such as ChIRP-seq and CHART-seq have been used to identify genome-wide binding sites of specific lncRNA to chromatin. However, these methods require prior knowledge of which particular lncRNAs are capable of binding to chromatin before ChIRP-seq or CHART-seq can be applied. Furthermore, ChIRP or CHART are limited to examining one chromatin associated RNA at a time. In this study, we describe a new technology, PIRCh-seq, which enables a global profiling of chromatin-associated RNAs through a robust method to crosslink endogenous RNA-chromatin interactions in living cells. Compared with current methods which predominantly detect nascent RNAs co-transcriptionally tethered to chromatin by RNA polymerase, PIRCh-seq significantly reduces the influence of nascent transcripts, and more clearly reveals relationships between chromatin-associated ncRNAs. Although the PIRCh approach cannot pinpoint the exact binding sites of the chromatin associated lncRNAs, and therefore does not inform whether each lncRNA is functioning in cis or trans, PIRCh is able to provide a significantly higher ratio of mature RNAs and thereby preserve the regulatory interactions in trans between lncRNAs and chromatin. Examples of some well-studied cases, such as *Xist*, *7sk*, *H19* and *KCNQ1OT1* etc. demonstrate that PIRCh-seq is likely generalizable to the majority of ncRNA. Additionally, PIRCh-seq identifies novel chromatin-associated lncRNAs and not only provides potential targets for mechanistic studies using ChIRP and CHART, but could also be extended to reveal the function and mechanisms of lncRNAs which are disease-relevant. However, the PIRCh-seq approach, like RIP/CLiP-seq like methods, may also be heavily contaminated with co-purified mRNA species that often compose more that 50% of RNA material. Therefore, further experimental and analytical improvements are required to truly capture chromatin-associated ncRNAs.

### PIRCh-seq classifies ncRNA putative function via histone modification and cell type-specific chromatin-RNA association

Another major advantage of the PIRCh method is that it utilizes antibodies to pull down chromatin with specific chemical modifications and thereby enables the classification of chromatin-associated ncRNAs with putative functions such as promoter, enhancer, silencer or bivalent. Since we performed PIRCh-seq with various histone modification antibodies and in different human and murine cell types, the dataset provides rich resources to study chromatin-associated ncRNAs in mammalian cells. In addition, different cell types and histone modifications did not show much technical variation, confirming that PIRCh-seq may be a useful technology to perform profiling of epigenetic-associated ncRNAs. Analogous to the types of gene regulatory elements bearing distinctive histone and DNA modifications, we developed a bioinformatics method to classify the putative biological functions of ncRNAs based on their enrichment patterns on chromatin with different histone modifications. Our method successfully arranged several well studied lncRNAs in the correct functional category, and predicted functions for hundreds of other ncRNAs from their chromatin association patterns. More importantly, when a similar analysis was performed on multiple cell types, chromatin state-specific ncRNA enrichment patterns were generally conserved, suggesting this is a reliable method for functional classification. Since ncRNA-chromatin interaction is likely a widespread epigenetic regulation mechanism in many cell types, our integrative approach in identifying and classifying chromatin-associated ncRNAs can be broadly applicable to many other cell types to deeper investigate ncRNA functions. However, chromatin association does not guarantee that a ncRNA will have a biological function; furthermore, the histone modification specific PIRCh-seq approach can only predict putative functions. As such, the true function of each ncRNA still requires further investigation beyond PIRCh-seq.

### RNA secondary structure affects RNA-chromatin interaction

We observed that RNAs attach to chromatin in a domain-specific manner. However, when we surveyed the enriched sites of chromatin-associated RNAs linked to various histone modifications in different cell types, we did not find significant sequence motifs, suggesting the existence of a complex mechanism responsible for the RNA-chromatin interaction. On the other hand, when we integrated PIRCh-seq signals with RNA structural information from previous icSHAPE and RNA modification m^6^A profiles and further evaluated structural information regarding the enriched domains, we found that ncRNAs were likely to bind to chromatin through single-stranded region or bases with m^6^A methylation. This may possibly be explained by the supposition that RNA-dependent recruitment of transcriptional activators and repressors may occur within a double-stranded structural region, and therefore, single-stranded regions are made more accessible to chromatin. In addition, chromatin interactions may also be allele-specific, especially when certain alleles result in distinct RNA secondary structures. In conclusion, when taken as a whole, these results open new avenues of inquiry and require further investigation to fully elucidate the molecular mechanisms of ncRNA-chromatin interaction.

## Author Contributions

KQ, QM, CC, HYC conceived the project. QM, CC, LL, PJB, KEMT, RL performed PIRCh-seq library generation and qPCR experiments. JF performed all data analysis with assistance from BH, PC, QM, JX and PD. KQ, HYC, JF, and QM wrote the manuscript with inputs from all authors.

## Data and code availability

The PIRCh-seq and ChIRP-seq data generated in this study can be obtained from NIH GEO with the accession number GSE119006 and is available by go to the following website https://www.ncbi.nlm.nih.gov/geo/query/acc.cgi?acc=GSE119006, and entering token “ulglmesubpmdbkx” in the box. Other published data sets used in this study are available and described in the Reporting Summary file. All in house developed codes/scripts were uploaded to Github website (https://github.com/QuKunLab/PIRCh).

## Acknowledgments

We thank the members of Chang and Qu labs for their discussions. This work was supported by the National Key R&D Program of China 2017YFA0102900 (to K.Q.), the National Natural Science Foundation of China grant (81788101, 91640113, 31771428 to K.Q.), the Chinese Government 1000 Youth Talent Program (to K.Q.), the research start-up from the University of Science and Technology of China (to K.Q.); the NIH grants P50-HG007735 and R01HG004361 (to. H.Y.C.); and the Stanford Dean’s Fellowship (to Q.M.). We thank the USTC supercomputing center and the School of Life Science Bioinformatics Center for providing supercomputing resources for this project. H.Y.C. is an Investigator of the Howard Hughes Medical Institute.

## Disclosure

H.Y.C. is affiliated with Accent Therapeutics (co-founder and advisor), 10X Genomics (advisor), and Spring Discovery (advisor).

## FIGURE LEGENDS

**Supplementary Figure 1.**
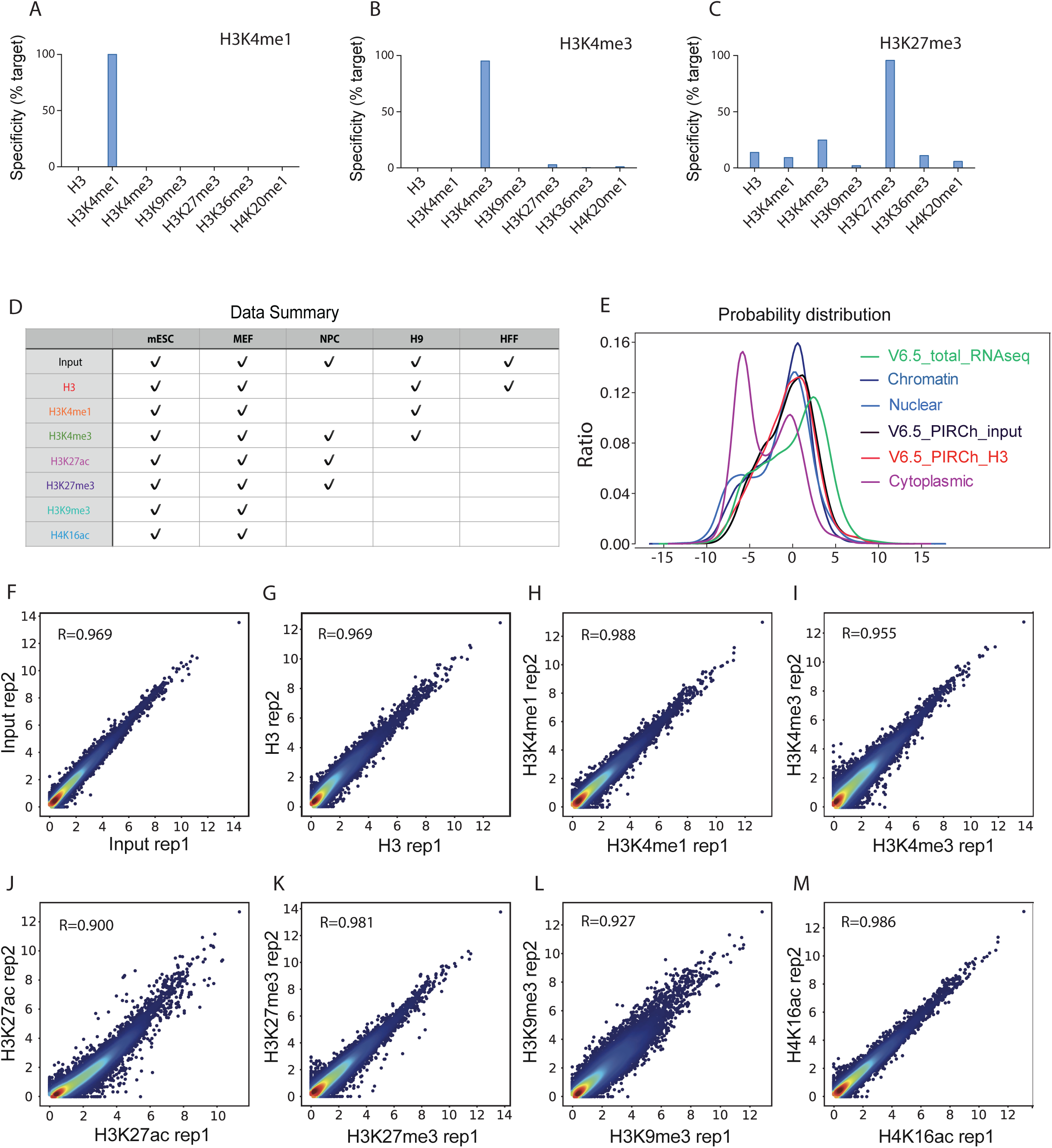
Quality control of histone modification specific PIRCh-seq experiments on distinct cell types. **A-C.** The specificity of IP of different antibodies H3K4me1 (A), H3K4me3 (B), and H3K27me3 (C) after glutaraldehyde crosslinking using modified mononucleosomes with barcodes. 7 different mononucleosomes with barcodes were tested. **D.** Table summarizing PIRCh-seq experiments performed in this paper. **E.** Kernel density estimation (KDE) plot of the gene expression from different subcellular RNA sequencing data and PIRCh-seq data. **F-M.** Scatter plots of expressed transcripts (log2) in two PIRCh-seq replicates with correlation score R on different histone modification, H3, H3K4me1, H3K4me3, H3K27ac, H3K27me3, H3K9me3, and H4K16ac respectively.

**Supplementary Figure 2.**
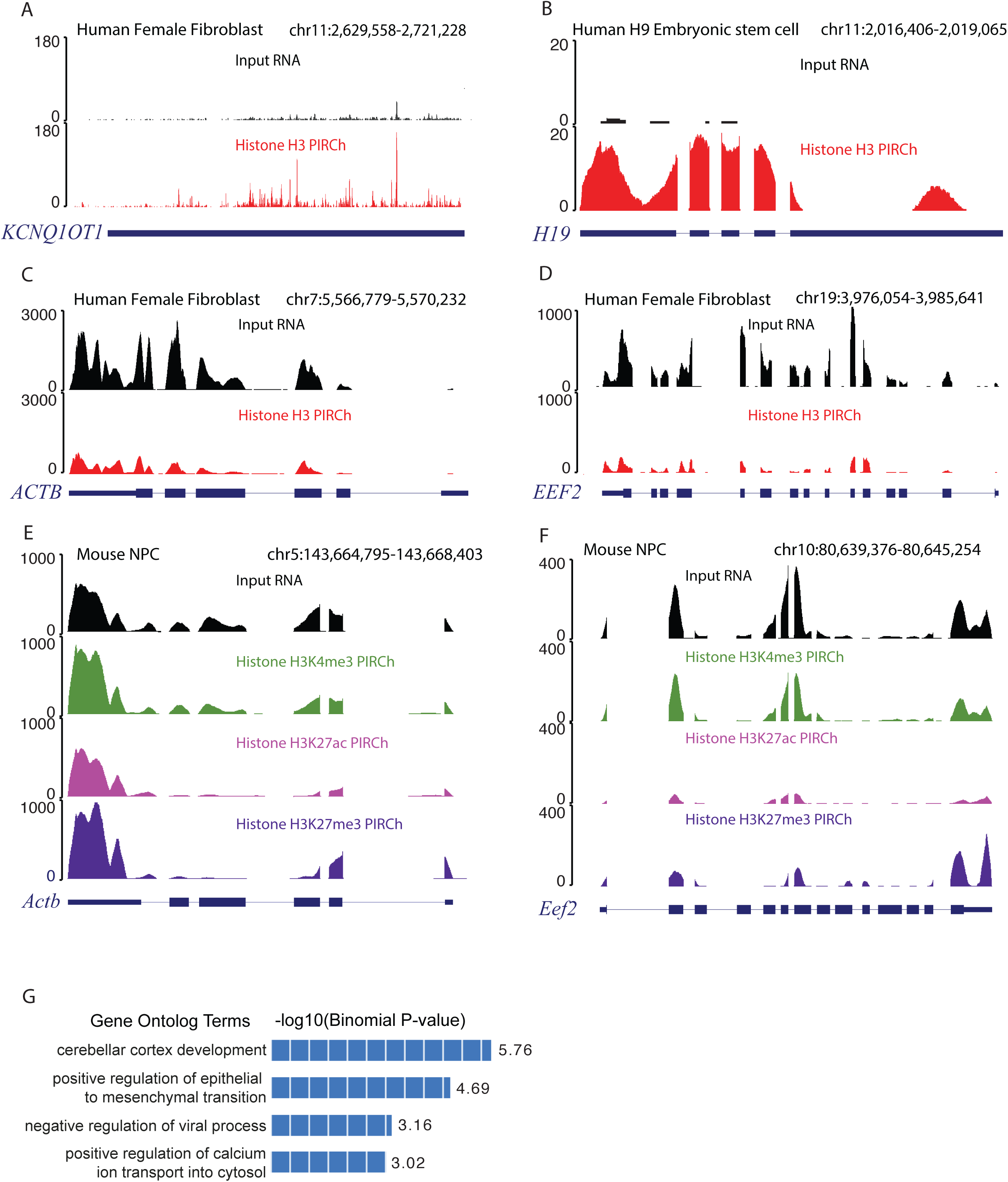
PIRCh-seq effectively and finely enriches RNA associated with chromatin. **A-B.** Normalized UCSC tracks of input and histone H3 PIRCh-seq signals on lncRNA *KCNQ1OT1*(A) in human female fibroblast cells and *H19* (B) in human H9 embryonic stem cells. **C-D.** Normalized UCSC tracks of input and histone H3 PIRCh-seq signals on protein coding genes *ACTB* (C), and *EEF2* (D) in human female fibroblast cells. **E-F.** Normalized UCSC tracks of input and histone modification specific PIRCh-seq signals on protein coding gene *Actb* (E) and *Eef2* (F) in mouse neuronal precursor cells. **G.** Top 5 enriched gene ontology of the *lnc-Nr2f1* ChIRP-seq peaks using GREAT.

**Supplementary Figure 3.**
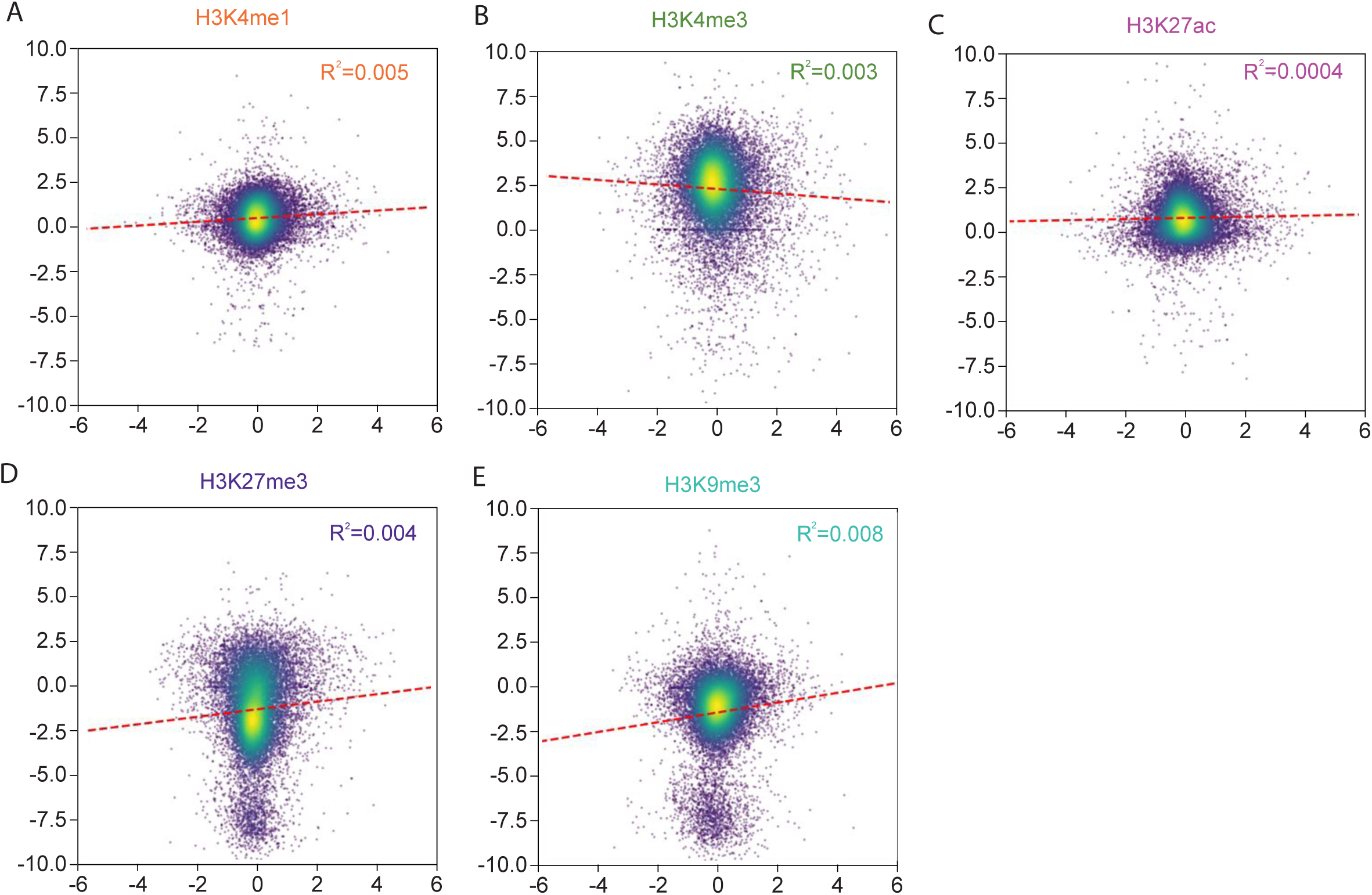
PIRCh-seq captures low nascent transcription. **A-E.** Scatter plot of the PIRCh-seq (y-axis) signal over input vs the corresponding ChIP-seq (x-axis) signal over input for all the expressed genes in mESC, with linear regression (red dotted line). Colors represent the density of point.

**Supplementary Figure 4.**
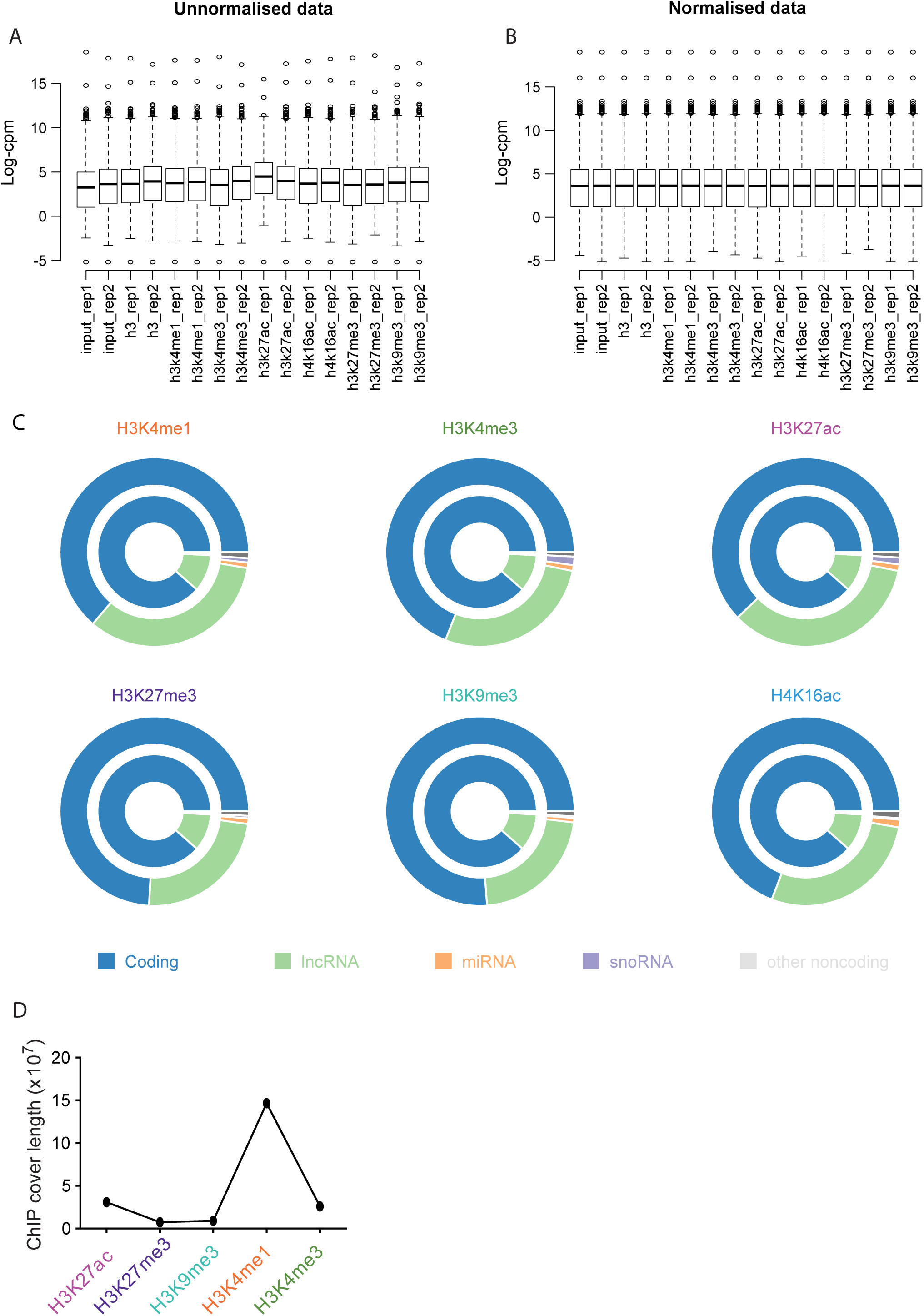
ncRNAs are more enriched on chromatin than protein coding genes. **A-B.** Box-plots of the PIRCh-seq signal before (A) and after (B) normalization using the limma algorithm in R. cpm represents count per million, and log scale is shown. Center lines represent mean values; box limits represent the interquartile range; whiskers each extend 1.5 times the interquartile range; dots represent outliers. **C.** Circle plots showing the distribution of the expressed (inner circle) and PIRCh enriched (outer circle) RNA types associated with different histone modifications. ncRNAs are highly enriched in PIRCh compared with coding genes. **D.** Total length of the genomic regions (in bp) covered by each histone modification ChIP-seq peak in mESC.

**Supplementary Figure 5.**
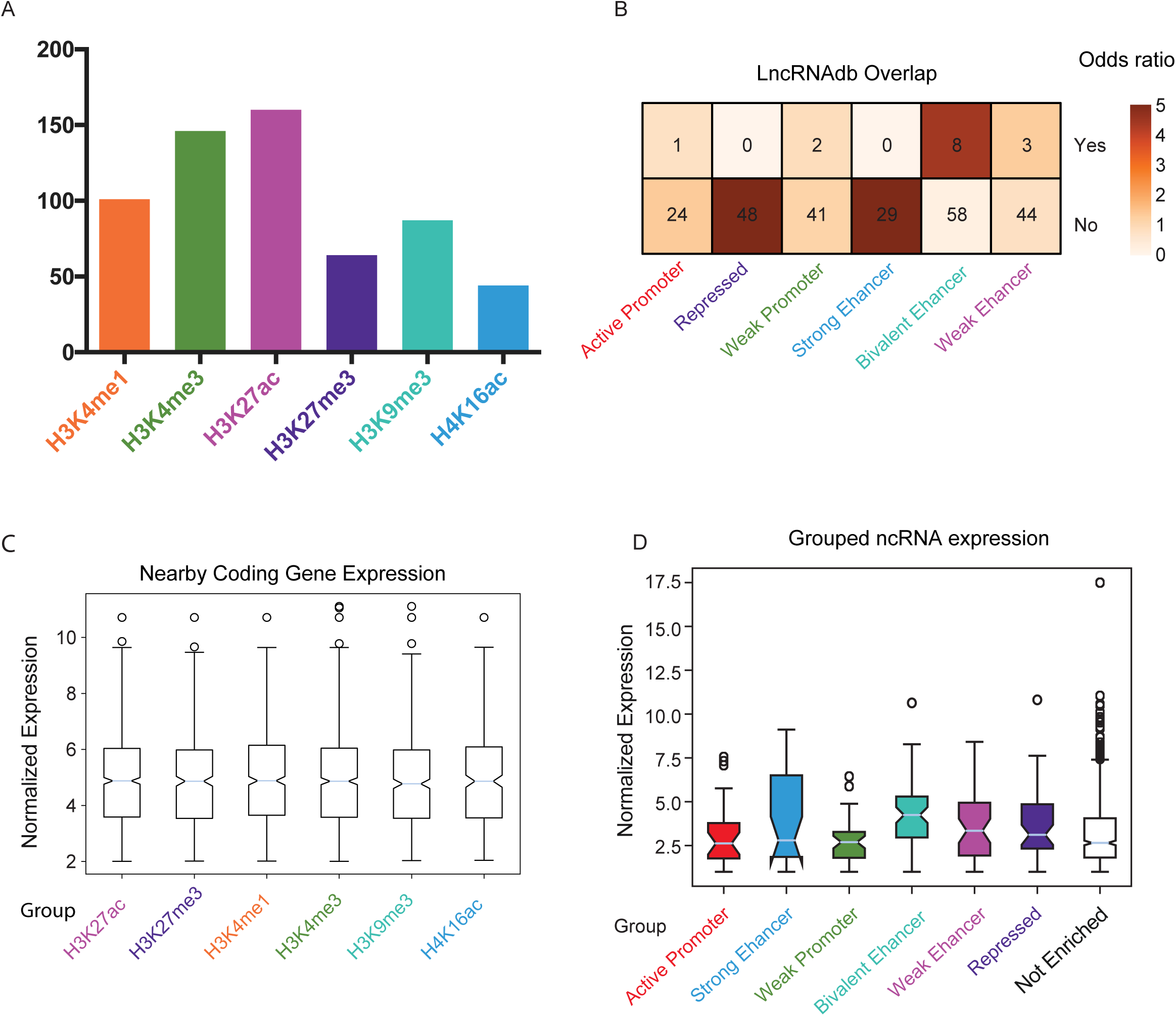
The chromatin-RNA association of ncRNAs give a hint of *cis* regulation. **A.** Bar chart showing the number of ncRNAs enriched at chromatin with specific histone modifications in mESC. **B.** The odds ratio of the PIRCh enriched ncRNAs overlap with the chromatin enriched ncRNAs defined in lncRNAdb. “Yes” means the ncRNA is both PIRCh enriched and found in lncRNAdb, and “No” means PIRCh enriched but was not identified in lncRNAdb. **C.** Box-plot of the expression of the coding genes near (+/-100Kb) each group of histone modification specific PIRCh-seq enriched ncRNAs. Center lines represent mean values; box limits represent the interquartile range; whiskers each extend 1.5 times the interquartile range; dots represent outliers. The expression of the coding genes that close to the ncRNAs enriched on active chromatin shows no significant difference between that with repressed chromatin. **D.** Box-plot of the expression of each groups of PIRCh cluttered ncRNAs defined in Figure 3D. Center lines represent mean values; box limits represent the interquartile range; whiskers each extend 1.5 times the interquartile range; dots represent outliers.

**Supplementary Figure 6.**
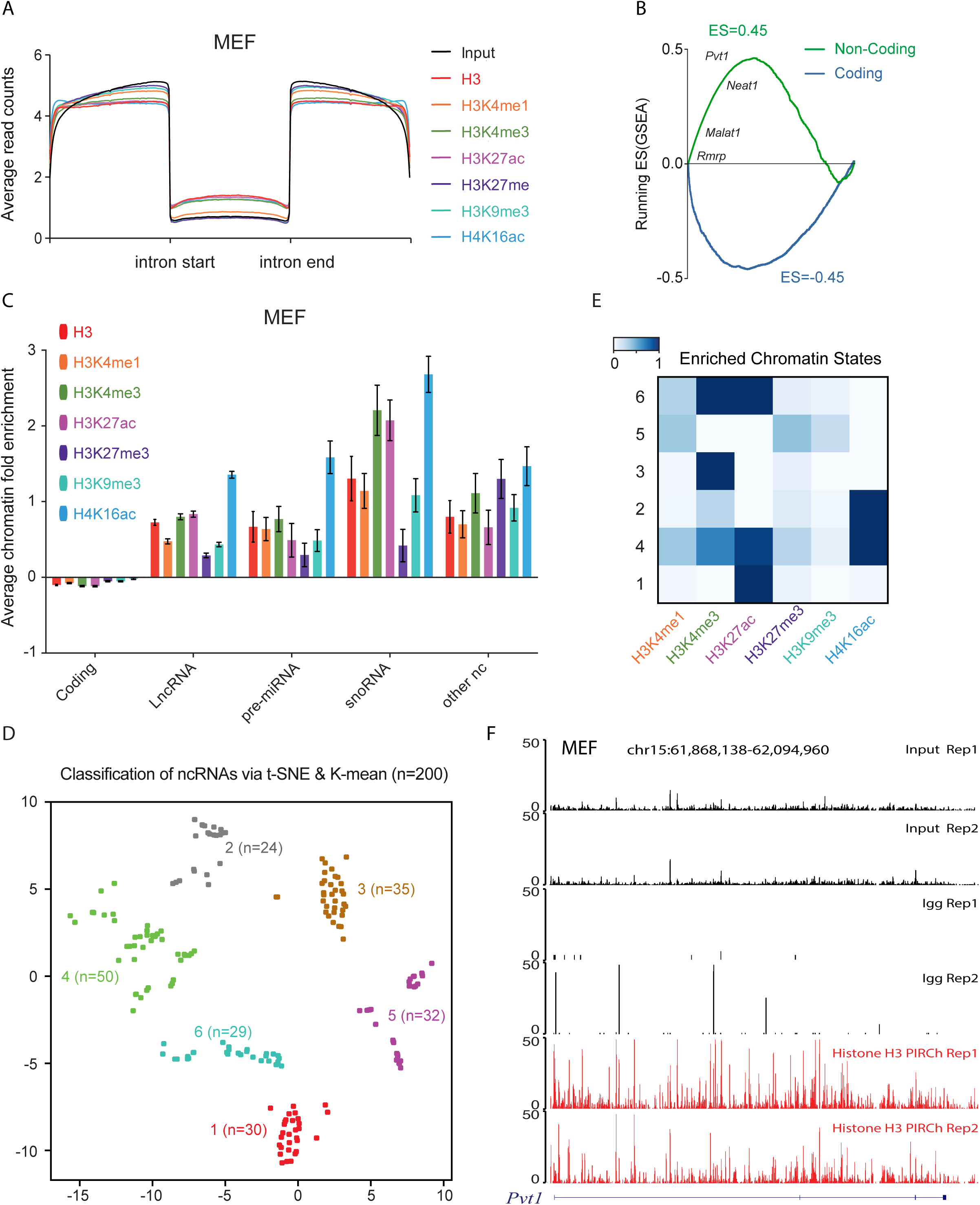
Pattern of ncRNA chromatin association is generally conserved in distinct cell types. **A.** Normalized average read coverage around introns from histone modification specific PIRCh-seq profiles (colored) and inputs (black) in MEF. **B.** Gene set enrichment analysis (GSEA) shows highly statistical enriched (FDR=0, P<0.0001) of non-coding genes (Green) and depleted of coding genes (Blue) on histone H3 in MEF. Genes were ranked by their histone H3 PIRCh enrichment scores. **C.** Average fold enrichment (calculated by limma in R) of the coding gene, lncRNA, pre-miRNA, snoRNA and other ncRNA from histone modification specific PIRCh-seq profiles (namely H3, H3K4me1, H3K4me3, H3K27ac, H3K27me3, H3K9me3, and H4K16ac) in MEF. Error bar shows the standard deviation from the mean. **D.** Functional classification of histone specific chromatin-RNA association patterns defined by chromHMM algorithm. **E.** Classification of the PIRCh-seq identified chromatin associated ncRNAs (n=200) in MEF. Scatter plot shows the t-SNE result on PIRCh-seq enrichment score matrix and annotated by K-means clustering. **F.** Normalized input, Igg and H3 PIRCh-seq profiles of lncRNA *Pvt1* in MEF.

**Supplementary Figure 7.**
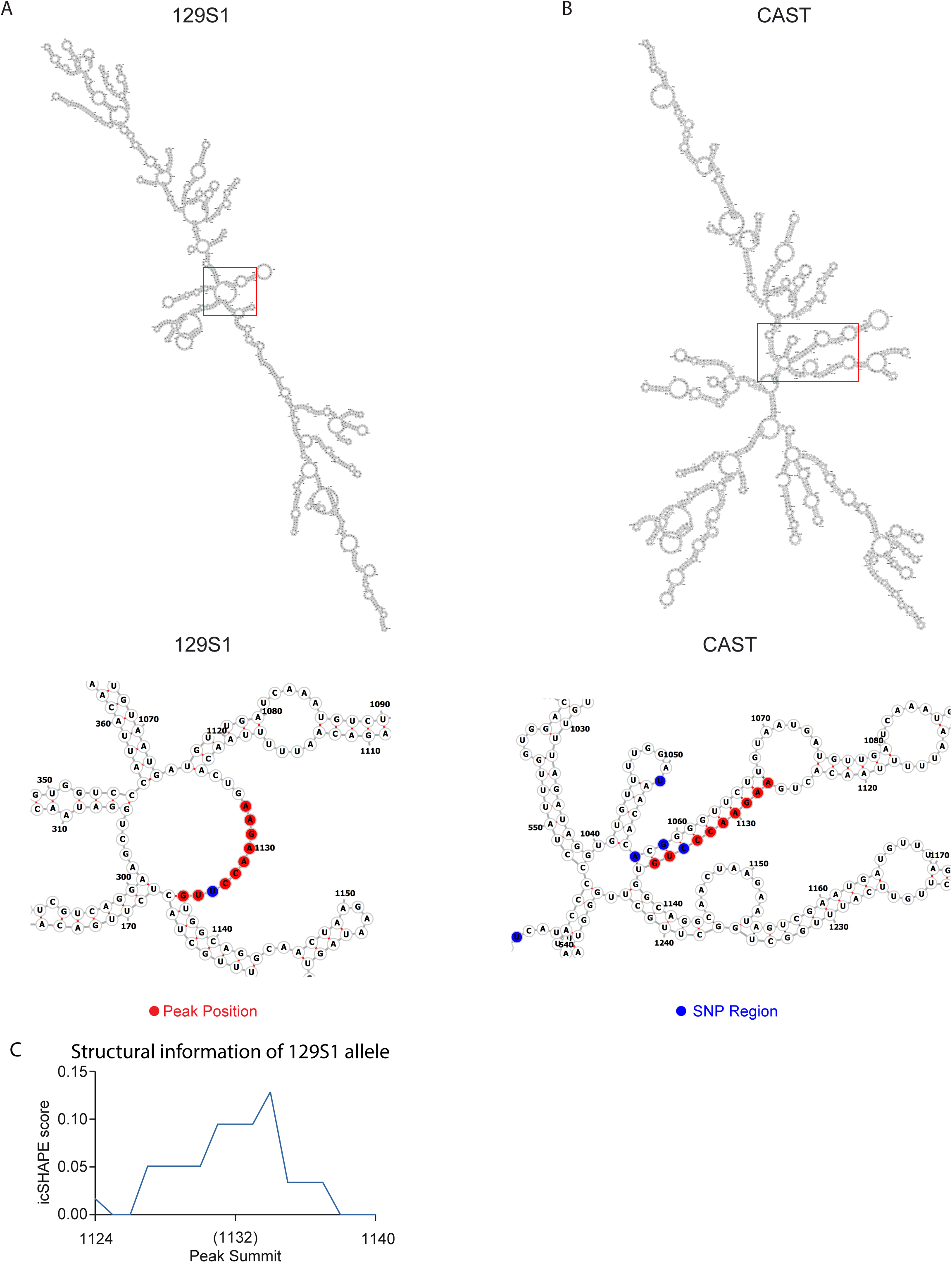
Allele specific RNA secondary structure and chromatin enrichment of lncRNA *Gas5*. **A-B.** RNAfold predicted the secondary structure of the 129S1 (A) and CAST (B) allele of *Gas5*. RiboSNithes are noted in blue and bases attached to chromatin (peaks) are shown in red. **C.** Structural information of 129S1 allele around the PIRCh enriched peak region. Data obtained from icSHAPE experiments on mESC.

**Supplementary Figure 8.**
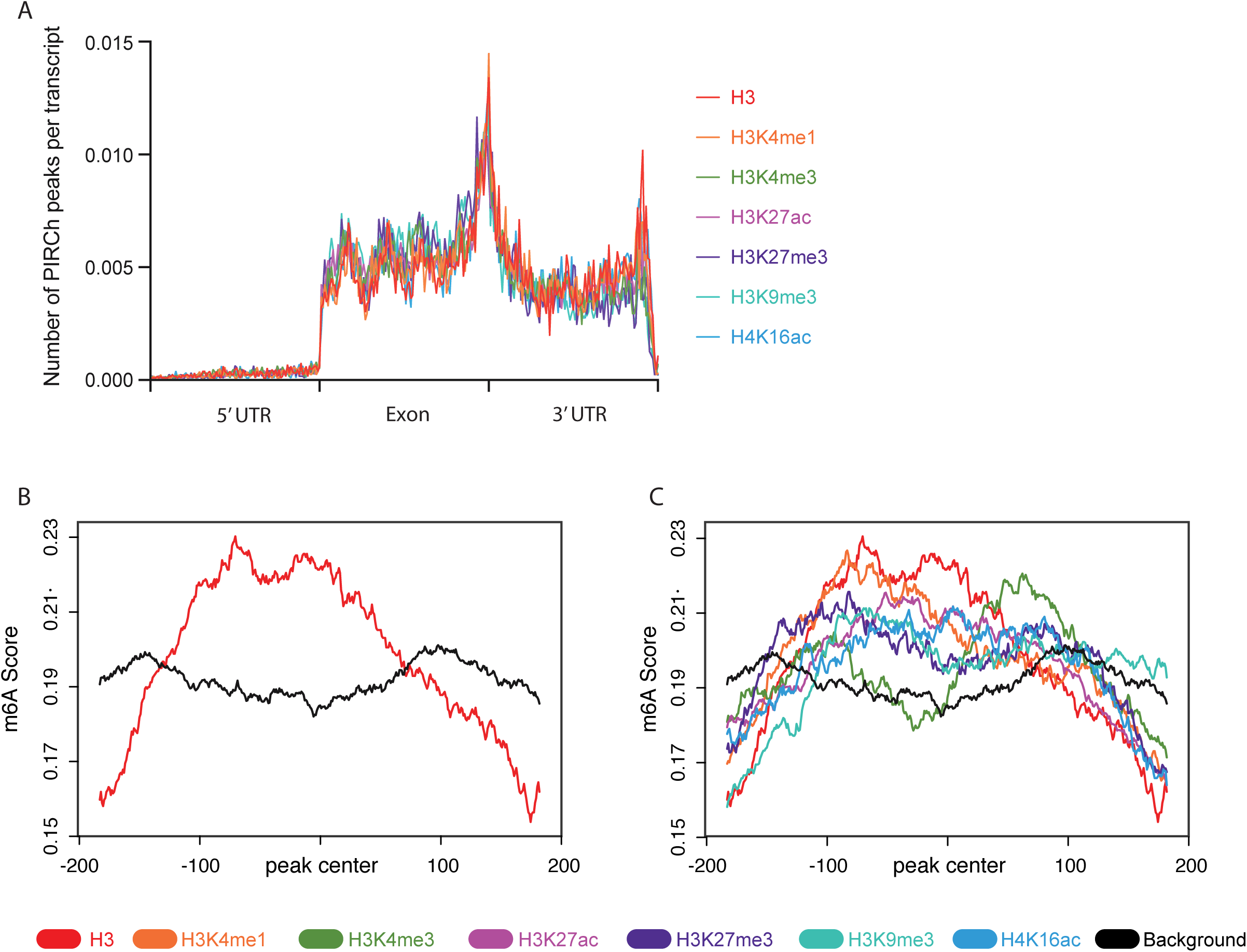
RNA m^6^A methylation affects chromatin-RNA association. **A.** Distribution of histone H3 and other chemical modification PIRCh-seq peaks along scaled transcripts. **B-C.** Average diagrams of m6A modification scores around bases attached to chromatin (peaks) from histone H3 (B) and all chemical modification specific (C) PIRCh-seq profiles, versus that from randomly selected background.

**Supplementary Table 1 Chromatin enriched ncRNAs in all mouse samples.**

**Supplementary Table 2 Allele specific chromatin enrichment in NPC.**

## METHODS

### Cell culture

V6.5 mouse ES cells were cultured on 0.2% gelatin-coated plates at 37°C with mES media: 500ml Knockout DMEM (Gibco), 90ml FBS, 6ml non-essential amino acid (NEAA, 100x, Gibco), 6ml glutamine or glutamax (200mM stock solution), 6ml Pen/Strep, 1ml BME and 60µlLIF (Millipore, ESG1106). Mouse embryonic fibroblast (MEF) cells were cultured at 37 °C and 5% CO2 in: 450ml DMEM, 50ml FBS, 5ml Pen/Strep, 5ml NEAA, 5ml pyruvate, 4ul beta-Mercaptoethanol. Mouse Neural Precursor cells (NPCs) were cultured in N2B27 medium (DMEM/F12 (Invitrogen, 11320-033), Neurobasal (Gibco, 21103-049), NDiff Neuro-2 Medium Supplement (Millipore, SCM012), B27 Supplement (Gibco, 17504-044)) supplemented with EGF and FGF (10 ng/ml, each) (315-09 and 100-18B, Peprotech). Cells were passaged using Accutase (SCR005, Millipore) and cultured on 0.2% gelatin-coated plates. H9 human embryonic stem cells were seeded in a feeder-free system using Matrigel hESC-Qualified Matrix (*354277*, Corning) and were maintained in Essential 8 media (A1517001, Thermo Fisher Scientific) as described previously^1^. Cells were passaged every three days as clumps with 0.5mM EDTA^1^. Human Female Fibroblasts (HFF) were cultured at 37 °C and 5% CO2 in DMEM supplemented with 1% pen/strep and 10% FBS.

### PIRCh-seq library preparation

To harvest the cells for PIRCh-seq, approximately 4×10^7^ cells were trypsinized and pooled into a 50ml falcon tube, after washing with 40ml of cold PBS once. Fresh 1% glutaraldehyde in room temperature PBS was created from 25% stock and remaining stock was discarded. The cell pellet was resuspended in 1ml of glutaraldehyde solution and a p1000 pipette was used to resuspend cells, and to top up to 40ml (1ml 1% glutaraldehyde / 1 million cells). After inverting several times, the tube was gently shaken for 10 minutes, and then quenched with 1/10 volume of 1.25M glycine. The tube was inverted several times, shaken gently for 5 min, and spun down 2000g for 4 min. The pellet was then washed once with 40ml cold PBS. The pellet was responded in 1ml/20million cells of cold PBS. Cells were aliquoted at 1ml each to a fresh eppendorf tube, and spun down 2000g for 4 min. After, supernatant was carefully aspirated, cell pellets were flash frozen, and stored at −80°C if necessary.

For sonication, prepared cell pellets were spun down at 2000g for 4 min and any remaining PBS was removed. Lysis per 20 million cells was performed with 1ml of lysis buffer (1% SDS, 50mM Tris 7.0, 10mM EDTA, 1mM PMSF, 0.1U/ul Superase-in (Ambion), 1x Proteinase inhibitor (Roche)). Lysate was then sonicated till the chromatin size was ∼300-2000bp and the lysate was clear. The lysates were spun down at 16000g for 10 min. Supernatants were flash frozen and stored at −80°C if necessary.

For PIRCh-seq library construction, chromatin was thawed and 10ul was taken as input. 200ul were aliquoted per reaction, and 400ul dilution buffer was added to each reaction. H3 or a specific histone modification antibody was then added (Dilution buffer: 0.01% SDS, 1.1% Triton X 100, 1.2 mM EDTA, 16.7 mM Tris 7.0, 167 mM NaCl, 1mM PMSF, 0.1U/ul Superase-in (Ambion), 1x Proteinase inhibitor (Roche)). The reaction was shaken end-to-end at 4°C overnight. 50ul Protein A dynabeads was used per 5ug antibody IP. Beads were washed with 5 times the original volume of dilution buffer 4 times. Notice that it is important not to exceed 200ul original volume of beads per tube. During the last wash, beads were aliquoted to 1 tube per reaction. The buffer was aspirated and, 200ul of the IP sample was used to resuspend and transfer beads to the IP sample. The reaction was shaken end-to-end at room temperature for 2 hours. The beads were then washed with 1ml wash buffer 4 times, and resuspended in 50ul IP elution buffer (1% SDS, 50mM NaHCO3). The reaction was then vortexed at setting 1 for 15 min. The supernatant was then transferred to a fresh tube and the bead elution was repeated. The supernatant was combined for a total of 100ul. 5ul 3M NaOAc was immediately added to neutralize pH. 10ul TurboDnase buffer and 1ul TurboDnase (Ambion) were added and the reaction was incubated 37°C for 30min. 3ul 500mM EDTA was added to eliminate divalent ions. 5ul Proteinase K (Ambion) was added, and the reaction was incubated at 50°C for 45 min.

To make our sequencing libraries, we extracted RNA using Trizol/chloroform, and precipitated the RNA with an equal volume of isopropanol. RNA pellet was washed in 1ml 70% EtOH, and pellets were resuspended in 10ul H2O. 1ul TurboDnase buffer was added, followed by 1ul TurboDnase, and the reaction at 37°C for 30min. 1.2ul of TurboDnase inactivating reagents were added. The reaction was vortexed for 3 minutes and spun down. The 10ul supernatant was heated at 75°C for 10 minutes to kill DNase. The reaction was purified using a Nugen Ovation v2 kit and eluted in 5uL for library preparation.

### ChIRP-seq library preparation

To determine the genome-wide localization of *lnc-Nr2f1*1 we followed protocols previously described^2^. ChIRP was performed using biotinylated probes designed against mouse lnc-Nr2f1 using the ChIRP probes designer (Biosearch Technologies). Independent even and odd probe pools were used to ensure lncRNA-specific retrieval as protocols previously described3. “Even” and “odd” sets of probes shared no overlapping sequences, as we performed two independent ChIRP-seq experiments with these two sets of probes separately. Two sets of data were then combined for downstream analysis (see below). Mouse NPC samples are crosslinked in 3% formaldehyde. RNase pre-treated samples are served as negative controls for probe-DNA hybridization. ChIRP libraries are constructed using the NEBNext DNA library preparation kit (New England Biolabs).

Sequencing libraries were barcoded using TruSeq adapters and sequenced on HiSeq or NextSeq instruments (Illumina).

### Experimental validation of antibody specificity after glutaraldehyde crosslinking using modified mononucleosomes with barcodes

To ensure that chemical crosslinking with glutaraldehyde did not affect antibody specificity, we followed previous study to test antibody specificity using SNAP-ChIP^4^. During IP pulldown, 15 uL of recombinant nucleosomes (SNAP-ChIP, EpiCypher, 19-1001) were fixed with fresh 1% glutaraldehyde. 1% glutaraldehyde was prepared on the same day in room temperature PBS from 25% stock. Fixation was performed for 10 minutes at room temperature with gentle shaking. The reaction was then quenched with 1/10 of the original reaction volume of 2.5 M glycine. Tubes were then inverted several times and incubated for 5 minutes at room temperature with gentle shaking.

500 uL of fixed chromatin were then added to each tube and pipetted up and down several times to mix well. 10 uL of nucleosomes mixed with chromatin were taken out of each tube to be used as input during the qPCR. One tablet of Roche cOmplete protease inhibitor was dissolved (Roche, 11697498001) in 50 mL of DI water to obtain a working solution of 50x protease inhibitor cocktail. 60 uL of 50x protease inhibitor was added to 3mL of blank dilution buffer (0.01% SDS, 1.1% Triton X100, 1.2 mM EDTA, 16.7 mM Tris pH 7.0, 167 mM NaCl). 1 mL of dilution buffer with protease inhibitor was then added to each reaction. 5 ug of appropriate detection antibody for IP pulldown was added to 300 uL of chromatin mixed with crosslinked nucleosomes for each condition. Samples were then incubated at 4°C overnight with end-to-end shaking.

IP product was eluted as specified during PIRCH library construction. DNA of interest was purified using a Zymo DNA Clean and Concentrator-5 kit (Zymo Research, D4013). The qPCR reaction was performed using Roche’s LightCycler and Brilliant II SYBR® Green QRT-PCR Master Mix (Agilent). We analyzed enrichment for target histone modifications by amplifying unique DNA barcodes at the 3’ end, using primer sequences provided by EpiCypher.

### RT-qPCR

For qRT-PCR analysis, we used Roche’s LightCycler and Brilliant II SYBR® Green QRT-PCR Master Mix (Agilent).

### PIRCh-seq data alignment

Raw reads were uniquely mapped to mm9/hg19 using Tophat with default parameters^5^. Samtools and BedTools were used to transform the mapped bam file into bedGraph and bigwig files for visualization on the UCSC genome browser^6, 7^. RPKM and raw reads count for each gene were calculated by self-designed scripts with ensemble annotation, Homo_sapiens.GRCh37.75.gtf for human and Mus_musculus.NCBIM37.67.gtf, and a number of previous publications for mouse samples respectively^8^.

### Calculate exon/intron ratio to estimate nascent transcripts

To compare the exon/intron ratios between the PIRCh-seq profiles and other chromatin associated RNA detection technologies, we aligned raw reads to the same hg19 genome index with Tophat and calculated the reads mapped to intron/exon with ensemble annotation gtf file as described above5. For the average read counts around introns, three steps were taken: (1) scaled every intron based on its length, and extended 1 intron length up and down stream of the selected intron; (2) divided the entire region to 300 windows, and calculate the average number of read counts mapped in each window and then take log2 to scale down the values to avoid interferences from the outliers; (3) take average for all the windows among all introns. To estimate the correlation between the histone modification-specific PIRCh-seq profile with its corresponding ChIP-seq signals, we obtained ChIP-seq profiles of each histone modification in mESC from ENCODE. And then, for each expressed gene in mESC, the histone modification ChIP-seq signal over input on the gene exon were calculated as the ChIP signal for that gene, and were compared with the corresponding PRICh-seq enrichment score with the same histone modification, and our results indicated that there was no significant correlation with these two sets of signals.

### Gene set enrichment analysis (GSEA)

GSEA software was downloaded from (http://software.broadinstitute.org/gsea/index.jsp) at the Broad Institute website and was utilized to perform the significant differential chromatin enrichment from PIRCh-seq against ncRNA versus coding genes^9^. The ncRNA set was consist of the annotated snoRNA, snRNA, rRNA, lncRNA, miRNA and miscRNA.

### Data normalization and identification of the chromatin enriched RNAs

The chromatin enriched ncRNAs were identified through the limma algorism in R^10^. First, a data matrix was obtained, where each raw read was a gene and each column a sample, and the element of the matrix represented the number of raw reads from PIRCh-seq experiments and inputs. The values in this matrix were then normalized by the limma-voom method in R. After that, differential analysis was performed using the limma gene-wise linear model for each pair of PIRCh replicates over inputs. Non-coding RNAs with P-value<0.05 and log2 fold change over inputs>0 were defined as chromatin enriched. We obtained 258 chromatin enriched ncRNAs in mouse V6.5 cell line, 200 in MEF and 110 in NPC. Variation score of each gene was defined as the standard deviations of the fold change among all histone modification specific PIRCh-seq profiles. The Pearson correlation coefficients between each two PIRCh-seq experiments were calculated and unsupervised clustering of the correlation matrix was performed in Cluster.

### Computational validation of the PIRCh-seq enriched ncRNAs

In order to validate the PIRCh-enriched candidates by similar methods, we examined 96 published chromatin-association datasets from ChIRP/CHART/RAP/GRID-seq experiments collected by the LnChrom database^11^. We found a total of 23 expressed lncRNAs in the LnChrom database, including *Xist, Firre, Rmrp, Tug1* and etc., and all of them were positively enriched in our PIRCh experiment and 14 of which were significant with P-value<0.05, suggesting the high sensitivity of the PIRCh approach in identifying chromatin associated lncRNAs. Furthermore, we obtained the genomic binding sites (peaks) of 23 lncRNAs from the aforementioned experiments, and overlapped them with the histone ChIP-seq peaks^12^ and got a ratio of the overlap for each lncRNA. We then calculated the Spearman correlation coefficients of these ratios with their corresponding lncRNA’s PIRCh-seq enrichment scores in the same cell line (normalized by the total number of different ChIP-seq peaks), and found that these correlations were significantly higher than random permutations. Peak calling was performed by MACS213 with FDR<0.05.

### The chromatin association states of the enriched ncRNAs

To cluster chromatin-enriched ncRNAs in distinct groups for functional prediction, we performed t-SNE and K-mean clustering on the PIRCh enrichment score matrix with the chromatin associated ncRNAs. The proper K number (K=6) was determined by silhouette score^14^.

### Nearby coding gene expression comparison

To further evaluate the functional prediction for chromatin enriched ncRNAs, we first grouped chromatin enriched ncRNAs by functional classification, and then obtained lists of the nearby (+/-100Kb) coding genes. We then calculated the gene expressions of these coding genes and represented them in box-plots. Similarly, we obtained a different list of nearby coding genes if the chromatin enriched ncRNAs were classified based on their chromatin enrichment scores on each histone modification. The significance between each group was estimated by 2-tail Welch’s T-test.

### lnc-Nr2f1 ChIPR-seq analysis

To further validate the PIRCh-seq candidates, we performed ChIRP-seq on one of the H3K4me3 modified PIRCh-seq enriched lncRNAs named *lnc-Nr2f1*. Experimental methods were mentioned above, where independent “even” and “odd” probe sets were applied. LncRNA *lnc-Nr2f1* ChIRP-seq data were then analyzed by applying a previously published pipeline^3^, where the read alignment was performed in bowtie2 and peak calling in MACS2. Signals from even and odd ChIRP-seq profiles were then merged to reduce false positive caused by probes. We confirmed that *lnc-Nr2f1* associated genomic regions were indeed enriched with H3K4me3 but no other modifications in NPCs, where the NPC ChIP-seq data was obtained from GSE117289, indicating the high specificity of our PIRCh-seq approach.

### Allelic specific enrichment analysis in NPC

We first built the CAST/EiJ and 129S1/SvImJ reference genome. The vcf files containing the SNPs in the CAST and 129S1 strains were downloaded from the dbSNP database with the mm9 assembly^15^. Their corresponding genome fasta file was made by GATK toolkit FastaAlternateReferenceMaker and SelectVariants tools^16^. After that, the inputs and PIRCh-seq data in were re-aligned against the CAST and 129S1 indexes by TopHat2 with 0 mismatch (parameter -N 0) to improve the specificity5. The allele specific alignment files were then converted to the bedGraph and bigWigs format using BEDtools. For each gene, its allele specific expression and enrichment analysis was performed for every SNP on the list, and estimated the significance between CAST and 129S1 through the Mann-Whitney-Wilcoxon test, and P-value<0.05 was defined as significant.

### Enriched peak calling from PIRCh-seq profiles

To further investigate the underlying mechanism of RNA-chromatin association, we performed peak calling on PIRCh-seq profiles to identify the bases on each enriched RNA that were mostly affiliated with histone proteins. We first merged data from two replicates of each gene to minimize the experimental deviation bias, and smoothed the normalized read counts on each base through a 5bp sliding window, along with a 2bp step size. Peak calling was performed on the smoothed signal with a home-made script. We defined a peak in the local maximum that is 5-fold or more amplified relative to the median read counts of the transcript. Next, we applied a bootstrap method by randomly sampling 1000 times with reads from the transcripts, and then estimated the P-value of each peak as the percentage of cases that were more enriched than observed. Finally, we calculated the relative fold-change of each peak with respect to the input control. Significant peaks were filtered based on fold change and P-value. Finally, RNA structural and modification information was integrated with PIRCh-seq peaks for downstream analysis.

### icSHAPE analysis and structural prediction using RNAfold

To estimate the structure information around PIRCh peak, we integrate mouse V6.5 icSHAPE data from previous paper^17, 18^. Each transcript’s icSHAPE score was calculated by the original icSHAPE pipeline with default parameter. We used home-made script to count icSHAPE score around PIRCh peak(+/-200bp) among all transcripts, and the significance between histone-modification PIRCh peak and random background region was estimated by by 2-tail Welch’s T-test. In terms of *Gas5* in NPC, the structure information of 129S1 allele was represented by V6.5 icSHAPE data, since they have the same sequence. Structure prediction of 129S1 allele and CAST allele was performed by RNAfold web server with default parameter^19^. For 129S1 allele, the higher icSHAPE score at peak region indicate single strand structure, which is similar to the structure prediction from RNAfold. Furthermore, structure prediction of CAST allele of *Gas5* in NPC shows that riboSnitches around PIRCh peak might be the cause of the allele specific enrichment of *Gas5*’s in NPC.

### Statistics

For data presented in Figure 1B (RT-PCR), P-values were calculated via the Mann-Whitney-Wilcoxon test in Python. For data presented in Figure 2D & S6B (GSEA), enrichment score, P-values and FDR were calculated in GSEA. For data presented in Figure S2G, binomial P-values were calculated by GREAT. For all T-test presented in this paper, included Figure 2E, F, I, Figure 3F, Figure S5C, Figure 5D, E, F, G, P-values were calculated via two-tailed Welch’s T-test in Python. For data presented in Figure 4F, P-values were calculated via the Chi-square Test.

### Data integration

Mouse v6.5 ChIP-seq results were downloaded from GSE102518^12^. Mouse NPC ChIP-seq data were downloaded from GSE117289. Mouse 7SK ChIRP-seq results were downloaded from GSE69143^20^. Murine structural information and RNA modification information were collected from our previous publications^18, 21^. All RNA binding peaks in ChIRP/CHART/RAP/GRID-seq experiments were downloaded from LnChrom^11^. All mouse data was analyzed using the mm9 assembly and all human data using hg19 assembly.

